# Red blood cell dynamics during malaria infection challenge the assumptions of mathematical models of infection dynamics

**DOI:** 10.1101/2024.01.10.575051

**Authors:** Madeline A.E. Peters, Aaron A. King, Nina Wale

## Abstract

For decades, mathematical models have been used to understand the course and outcome of malaria infections (i.e., infection dynamics) and the evolutionary dynamics of the parasites that cause them. The extent to which this conclusion holds will in part depend on model assumptions about the host-mediated processes that regulate RBC availability, i.e., removal (clearance) of uninfected RBCs and supply of RBCs. Diverse mathematical functions have been used to describe host-mediated RBC supply and clearance in rodent malaria infections; however, the extent to which these functions adequately capture the dynamics of these processes has not been quantitatively interrogated, as *in vivo* data on these processes has been lacking. Here, we use a unique dataset, comprising time-series measurements of erythrocyte (i.e., mature RBC) and reticulocyte (i.e., newly supplied RBC) densities during *Plasmodium chabaudi* malaria infection, and a quantitative data-transformation scheme to elucidate whether RBC dynamics conform to common model assumptions. We found that RBC supply and clearance dynamics are not well described by mathematical functions commonly used to model these processes. Indeed, our results suggest said dynamics are not well described by a single-valued function at all. Furthermore, the temporal dynamics of both processes vary with parasite growth rate in a manner again not captured by existing models. Together, these finding suggest that new model formulations are required if we are to explain and ultimately predict the within-host population dynamics and evolution of malaria parasites.

## 1 INTRODUCTION

Red blood cells (RBCs) are the primary target cells of malaria parasites during the blood-stage of infection and their availability changes dramatically over an infection’s course (Lamb and Langhorne, 2008; Timms et al., 2001; Mackinnon and Read, 1999; Wale et al., 2017b). These changes are caused by parasite destruction of RBCs, as well as a suite of host responses (reviewed by Deroost et al., 2016), and can have profound consequences for both host health and parasite fitness, for example via the anemia characteristic of malaria infections. As such, quantitatively understanding the causes and consequences of RBC dynamics *in vivo* is essential if we are to explain the disease and dynamics of malaria infections.

Over the last three decades, mathematical models have been used to quantitatively understand the various mechanisms by which variation in RBC dynamics comes about and the consequences of this variation for the population dynamics and evolution of malaria parasites (reviewed in Khoury et al. (2018) and Tables 1, 2). For example, Jakeman et al. (1999) used a mathematical model to suggest that the anemia of malaria infections results primarily from destruction of uninfected RBCs, rather than from direct destruction by parasites or impaired RBC production (dyserythropoiesis). Others have elucidated the consequences of anemia for the population dynamics and evolution of malaria parasites. Theoretical studies have posited that anemia induces RBC limitation, resulting in the cessation of parasite growth and intense inter-parasite competition for RBCs (Haydon et al., 2003; McQueen and McKenzie, 2004; Mideo et al., 2008; Wale et al., 2019). Over evolutionary time, these competitive interactions are hypothesized to have driven selection for parasite traits that impact host health, such as the preference of different species for different ages of RBCs (“age preference”) and growth rate (De Roode et al., 2005; Antia et al., 2008; Pak et al., 2024).

**Table 1.** Functional forms commonly used to describe reticulocyte supply during murine malaria infections. Numbering corresponds to example functional forms displayed in Figure 1A. Lag refers to the amount of time in days between a given value of total red blood cell (RBC) density and the corresponding reticulocyte supply response. For “functional forms” that are non-autonomous, reticulocyte supply is an explicit function of time (i.e., does not depend on current or lagged RBC densities). Under compartmental aging, RBCs progress through age compartments and are removed from circulation upon leaving the final age compartment. *^∗^*RBC supply is not explicitly described.

|  | Reference | Functional form<br>(function of RBC deficit) | Lag (days) | Reticulocyte maturation<br>in circulation |
| --- | --- | --- | --- | --- |
| 1 | Anderson et al. (1989) | Constant | - | - |
| 2 | Antia et al. (2008) | Hill | 2.5 | Compartmental aging |
| 3 | Cromer et al. (2006) | Hill | 2 | 2-days |
| 4 | Gravenor et al. (1995) | Constant | - | - |
| 5 | Greischar et al. (2016) | Linear (decreasing) | 0 | - |
| 6 | Haydon et al. (2003) | Linear (decreasing) | 0 | - |
| 7 | Hellriegel (1992) | Constant | - | - |
| 8 | Hetzel and Anderson (1996) | Constant | - | - |
| 9 | Jakeman et al. (1999) | Non-autonomous;<br>constant; no supply | 0 | Compartmental aging |
| 10 | Kamiya et al. (2021) | Constant with additional<br>supply increasing with<br>RBC deficit | 2 | - |
| 11 | Lim et al. (2013) | Constant | - | Not applicable or<br>compartmental aging |
| 12 | McQueen and McKenzie (2004) | Non-autonomous | 0 | Compartmental aging<br>assuming 36 hours<br>total on average<br>with variance of<br>~8 hours |
| 13 | Metcalf et al. (2011) | Linear* | 0 | Inferred to be 3-days |
| 14 | Metcalf et al. (2012) | Non-autonomous* | 0 | - |
| 15 | Mideo et al. (2008) | Linear (decreasing) | 1-4 | 3-days |
| 16 | Miller et al. (2010) | Linear (decreasing) plus<br>upregulation based on<br>lagged RBC deficit | 0-6 | Estimated |
| 17 | Thakre et al. (2018) | Hill | 2 | Compartmental aging<br>with reticulocytes<br>maturing in 36 hours |
| 18 | Wale et al. (2019) | Non-autonomous,<br>non-parametric | - | 1-day |

**Table 2.** Functional forms commonly used to describe uninfected RBC (red blood cell) clearance during malaria infections. Numbering corresponds to example functional forms displayed in Figures 1B-C. For “functional forms” that are non-autonomous, uninfected RBC clearance is an explicit function of time (i.e., does not depend on RBC densities). *^∗^*RBC destruction is not explicitly described. *^∗∗^*Clearance occurs through aging out of the oldest RBC class.

| Reference | RBC clearance |
| --- | --- |
| 1 Anderson et al. (1989) | Increasing linear function of RBC density |
| 2 Antia et al. (2008) | Constant** |
| 3 Cromer et al. (2006) | Increasing linear function of RBC density, with an increased clearance rate after day 6 |
| 4 Gravenor et al. (1995) | Increasing linear function of RBC density |
| 5 Greischar et al. (2016) | Increasing linear function of RBC density |
| 6 Haydon et al. (2003) | Unspecified |
| 7 Hellriegel (1992) | Increasing linear function of RBC density |
| 8 Hetzel and Anderson (1996) | Increasing linear function of RBC density |
| 9 Jakeman et al. (1999) | Non-autonomous; senescence only |
| 10 Kamiya et al. (2021) | Constant baseline mortality, additional clearance scaling with parasite density |
| 11 Lim et al. (2013) | Increasing linear function of RBC density |
| 12 McQueen and McKenzie (2004) | Constant** |
| 13 Metcalf et al. (2011) | Linear* |
| 14 Metcalf et al. (2012) | Non-autonomous* |
| 15 Mideo et al. (2008) | Constant fraction |
| 16 Miller et al. (2010) | Increasing linear function of RBC density |
| 17 Thakre et al. (2018) | Increasing linear function of RBC density |
| 18 Wale et al. (2019) | Non-autonomous, non-parametric |

Critically, the robustness of model-derived inferences about the disease and dynamics of malaria infections depend on the validity of assumptions underlying those models (Childs and Buckee, 2015; Peters et al., 2021). In particular, assumptions about the processes by which RBCs are supplied and cleared will be especially important, since these processes are major drivers of the availability and age structure of RBCs. The majority of studies that include models of within-host rodent malaria dynamics describe RBC supply and uninfected RBC (uRBC) clearance as a constant and/or as a simple function of RBC abundance (Tables 1, 2; Figure 1). For example, the supply of immature RBCs (reticulocytes) is commonly modeled (12 of the 18 papers in Table 1) as a constant function of time (Anderson et al., 1989; Gravenor et al., 1995; Lim et al., 2013; Khoury et al., 2018), as an increasing linear function of the deficit in uninfected RBC density (a “homeostatic” model) (Greischar et al., 2016; Haydon et al., 2003; Mideo et al., 2008) or as a sigmoidal function of this uninfected RBC deficit (Antia et al., 2008; Cromer et al., 2006; Thakre et al., 2018). Although these assumptions are appealing in their simplicity and have been necessitated by the technical challenges of quantifying the dynamics of RBC supply and clearance *in vivo*, recent work implies that these simple functions may not best represent RBC dynamics. For example, a significant body of empirical work suggests that the immunohematological processes responsible for RBC supply change dramatically upon infection and may vary with parasite traits, counter to the “homeostatic” model of RBC supply (Ruan and Paulson, 2023; Lamb and Langhorne, 2008; Lin et al., 2017; Wale et al., 2017b).

**Figure 1.**
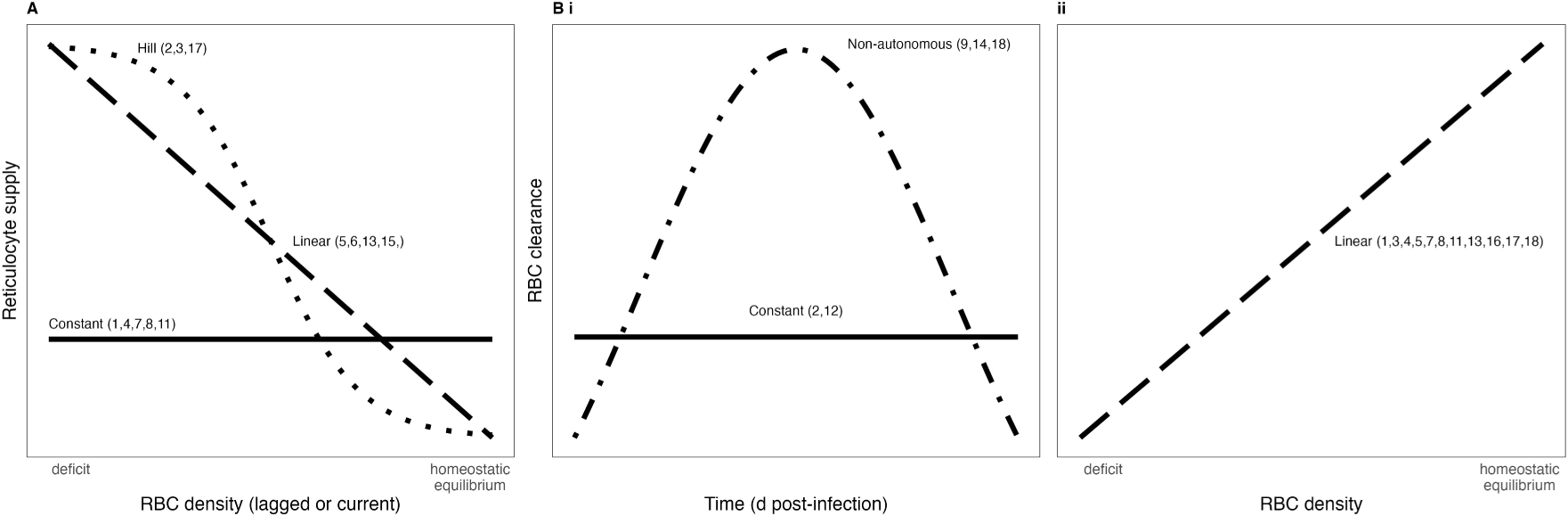
Functional forms commonly used to describe RBC supply and uninfected RBC clearance during malaria infections. Numbers associated with each functional form correspond to references in Tables 1, 2.

Here, we investigate the validity of assumptions about RBC supply and clearance commonly invoked in studies modelling within-host rodent malaria dynamics, using a unique dataset and data-transformation scheme that permit us to quantify these regulatory processes throughout the course of infection. Specifically, in addition to standard measures of RBC and parasite dynamics, our dataset includes quantitative measures of reticulocyte supply, which allow us to directly investigate how reticulocyte supply and uninfected RBC clearance change over the course of an infection (e.g., with time or RBC density). Importantly, our intent is not to discover the “correct” functional forms that describe RBC supply and clearance during acute malaria infection; rather we aim to examine whether commonly employed assumptions about RBC supply and clearance are sufficient to explain our data. We find that mathematical functions commonly used to describe RBC supply and clearance do not capture our data well. Rather, our results suggest we need new data streams and modeling strategies if we are to fully understand malaria infection dynamics and evolution.

## 2 MATERIALS AND METHODS

Here, we conduct a new analysis of a previously published dataset and data-transformation scheme to elucidate (i) to what extent the functions commonly used to describe red blood cell (RBC) supply and clearance describe our data and (ii) whether these functions change with an experimental manipulation known to change infection dynamics. Full details about the data-transformation scheme and the data can be found in Wale et al. (2019). We describe each briefly below.

### 2.1 Hosts and parasites

Hosts were 6- to 8-week old C57BL/6J female mice. 12 mice were infected with 10^6^ *Plasmodium chabaudi* parasites of a pyrimethamine-resistant strain denoted AS_124_.

To generate variation in the growth rate and dynamics of infection, we varied the supply of *para*-aminobenzoic acid (pABA) to mice. pABA is not a nutrient for the host (Fenton et al., 1950) but is nonetheless routinely supplemented to experimentally-infected mice (Gilks et al., 1989), as it significantly stimulates parasite growth rate (Hawking, 1954; Jacobs, 1964; Wale et al., 2017b). We have found that the dynamics of malaria infections, particularly those caused by pyrimethamine-resistant parasites (Wale et al., 2017a,b), varies with the concentration of pABA supplemented to mice. We thus manipulated pABA to generate variation in infection dynamics and to investigate the relationship between parasite growth rate and RBC dynamics, in particular. To emphasize that we are not performing a study of host nutrition, we refer to pABA as a “parasite nutrient”, throughout. In the experiment, groups of 3 mice each received either a 0.05% (high), 0.005% (medium), 0.0005% (low) or 0% (unsupplemented) solution of pABA as drinking water, initiated a week prior to parasite inoculation. Infections were monitored daily from day 0 (i.e., the day of inoculation) to day 20 post-inoculation. Following daily blood sampling from the tail, parasite densities, erythrocyte densities (i.e., mature RBC densities) and reticulocyte densities (i.e., immature RBC densities) were measured via quantitative PCR, Coulter counting and flow cytometry (Figure 2A, S1, *Supplementary Material*). One mouse in the high pABA treatment died on day 8. One mouse in each of the high and medium pABA treatments received fewer parasites than was intended; they were omitted from our analyses. The experiment was reviewed by the Institutional Animal Care and Use Committee of Pennsylvania State University, where the experiment was conducted (IACUC protocol 44512-1, P.I. Dr. A. Read).

**Figure 2.**
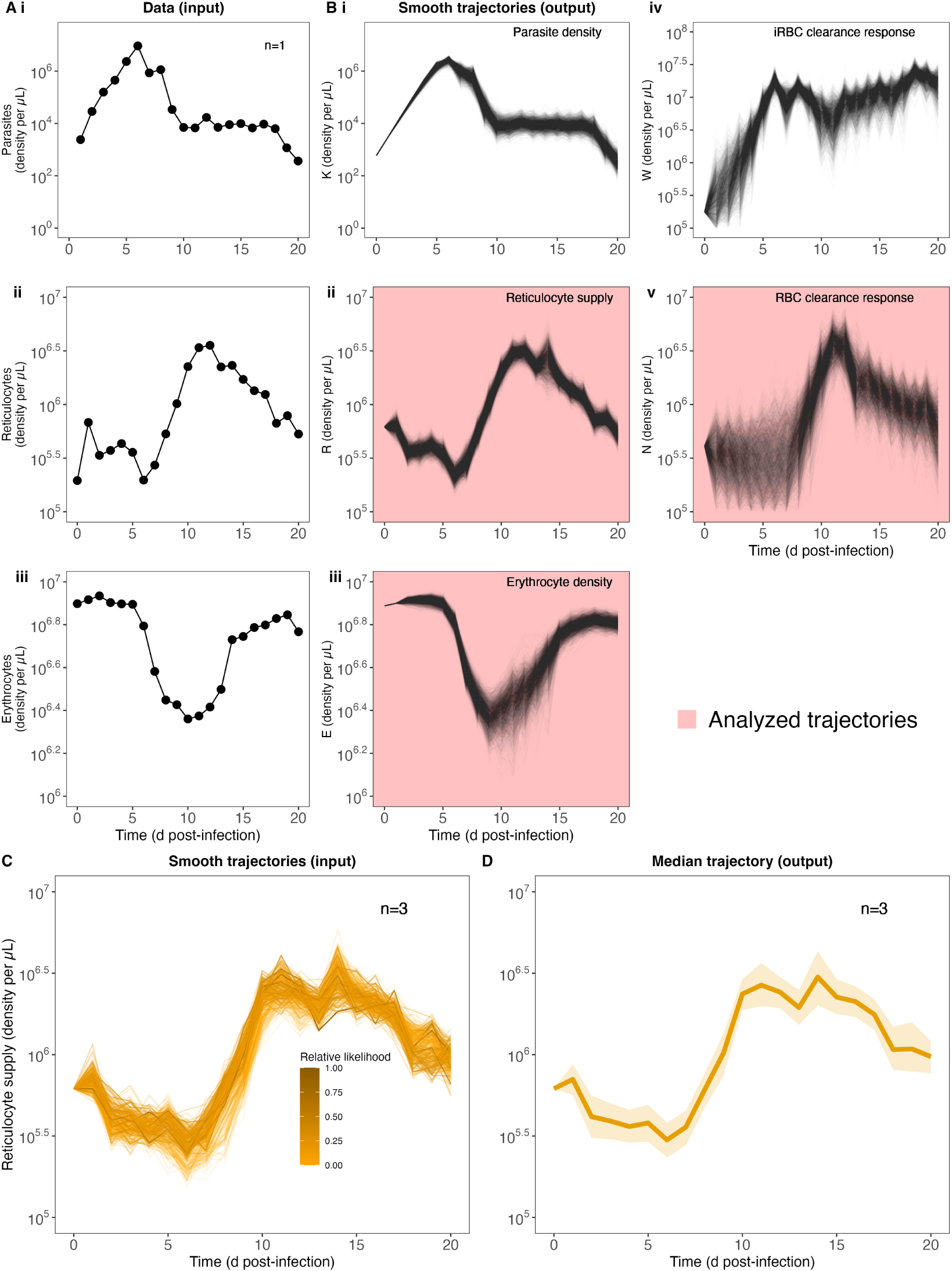
Quantifying RBC supply and clearance using a data-transformation scheme. Our strategy begins with raw data (A) of parasite, reticulocyte and erythrocyte density during infection (data from 1 mouse shown for illustration purposes, cf. Figure S1 (*Supplementary Material*) for full dataset). (B) Fitting these to the data transformation scheme yields 2000 trajectories of each of 5 different variables: parasite density *K*, reticulocyte density *R*, erythrocyte density *E*, targeted killing *W* and indiscriminate killing *N*. Note that variables *R*, *E* and *N* are used in the next step of the analysis. (C) To calculate a median trajectory of each of our three focal variables for each pABA treatment, we sampled from the trajectories of each mouse in each treatment (trajectory color/opacity indicates the likelihood). For ease of readability, we show only 300 trajectories from the unsupplemented (0%) pABA treatment (of 6000 total trajectories, with 2000 per mouse). This process yields a median (dark line, 90% confidence interval, shading) trajectory for each group of mice (unsupplemented pABA treatment shown in (D)).

### 2.2 Data transformation

To quantify the host RBC supply and clearance responses during infection, we used the approach first presented by Wale et al. (2019). Notably, rather than generating quantitative *predictions* of malaria infection dynamics (*a la* the references in Tables 1, 2), this approach transforms three data streams (densities of erythrocytes, reticulocytes, parasites; Figure 2A) into three host responses, which together determine the supply and clearance of RBCs (Figures 2Bii,iv,v). We thus refer to this approach as a “data-transformation scheme”, as opposed to a model.

The equations posit that

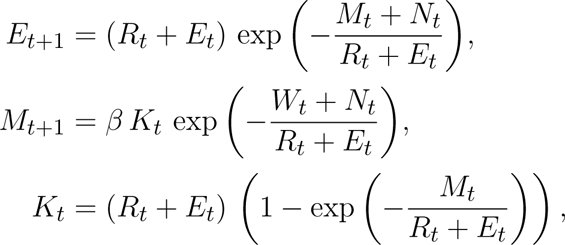

where, on day *t* post-infection, *E_t_*and *R_t_*are unparasitized erythrocyte and reticulocyte densities, respectively, *M_t_*is merozoite density and *K_t_*is the density of parasitized RBCs. These equations express the assumptions that (i) reticulocytes entering circulation on day *t* mature to become erythrocytes on day *t* + 1 (Ney, 2011; Gronowicz et al., 1984; Koury et al., 2005; Wiczling and Krzyzanski, 2008; Noble et al., 1989), (ii) *P. chabaudi* merozoites attack unparasitized RBCs without respect to age (Jarra and Brown (1989); Yap and Stevenson (1994); Carter and Walliker (1975); Taylor-Robinson and Phillips (1994), but see Antia et al. (2008); Mideo et al. (2008)), (iii) parasitized RBCs burst in one day to produce *β* merozoites on average, (iv) infected RBCs produce the same number of merozoites regardless of RBC age, (v) burst size is independent of the number of infecting merozoites, (vi) multiply-infected RBCs have the same chance of survival as singly infected RBCs, (vii) parasitized RBCs are removed by the immune system at a rate dependent on the time-varying quantity *W_t_*, and (viii) irrespective of their age and parasitization, RBCs are removed from circulation at a rate dependent on the time-varying quantity *N_t_*. All model terms are described in Table 3.

**Table 3.**
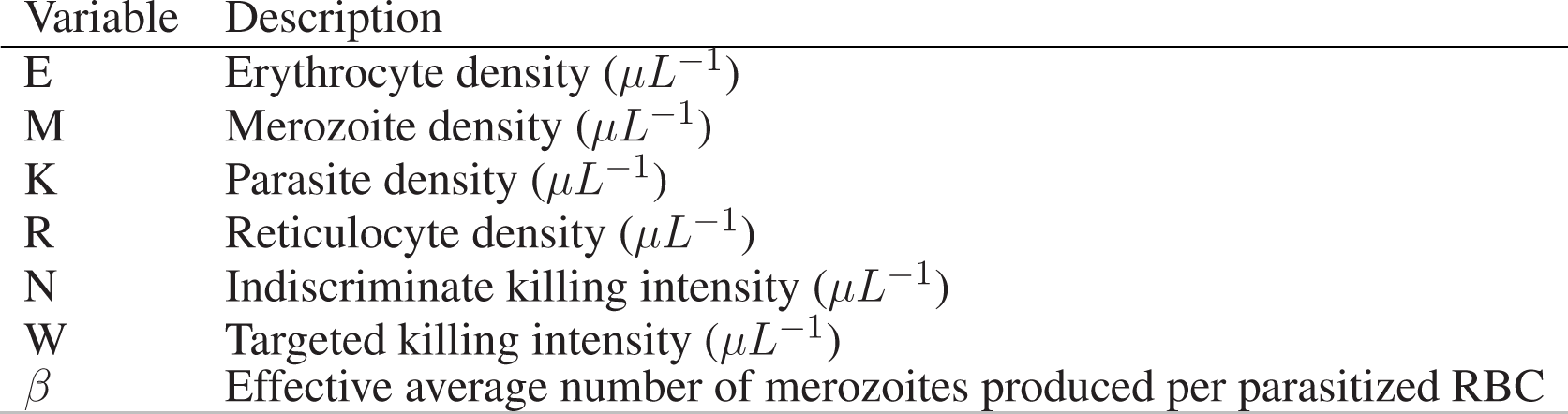
Data-transformation scheme terms and parameters. Indiscriminate killing refers to the immune killing of both uninfected and infected red blood cells (RBCs), while targeted killing refers to the immune killing of only infected RBCs. Note that *N* and *W* are scaled to be on the same scale as *R*.

| Variable | Description |
| --- | --- |
| $E$ | Erythrocyte density ($\mu L^{-1}$) |
| $M$ | Merozoite density ($\mu L^{-1}$) |
| $K$ | Parasite density ($\mu L^{-1}$) |
| $R$ | Reticulocyte density ($\mu L^{-1}$) |
| $N$ | Indiscriminate killing intensity ($\mu L^{-1}$) |
| $W$ | Targeted killing intensity ($\mu L^{-1}$) |
| $\beta$ | Effective average number of merozoites produced per parasitized RBC |

We estimate the values of the state variables (*R_t_*, *W_t_*, *N_t_*,) and the *β* parameter from the data, as follows. The transformation is regularized by assuming that the time-dependent functions log *R_t_*, log *W_t_*and log *N_t_* are Gaussian Markov random fields (GMRF), each with its own intensity, and that the measured values of reticulocyte, RBC and parasitized cell densities on day *t* are log-normally distributed about *R_t_*, *R_t_* + *E_t_* and *K_t_*, respectively. The GMRF intensities, error standard deviations and initial conditions were estimated from the data using iterated filtering using the **R** package **pomp** (King et al., 2016; Ionides et al., 2015). 2000 samples from the smoothing distribution for each of these variables were drawn for each individual infected mouse (Figure 2B). We further estimated *β* values for each pABA treatment using linear regression applied to the first 4 days of data (i.e., we take the slope of the curve for each treatment; Figure S2). Full details of these computations are provided in the original paper and its supplement (Wale et al., 2019); the code for the computations in this paper are available on GitHub at https://github.com/kingaa/malaria-rbc-dynamics and will be archived on Zotero upon acceptance of this paper.

### 2.3 Estimating clearance rate

In the majority of modeling studies, clearance rate is defined as the rate at which uninfected RBCs (uRBCs) are removed. Accordingly, we use the output of our data-transformation scheme to estimate 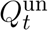, the probability on a given day *t* that an individual uRBC will be cleared by indiscriminate killing. We first define 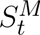, the probability that an RBC escaped infection by a malaria parasite and 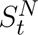, the probability that an RBC is not cleared by indiscriminate killing as:

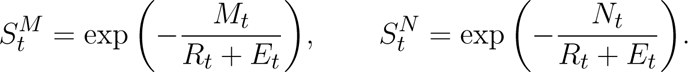

The probability that an uninfected RBC is cleared by indiscriminate killing on day *t* is then

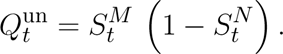

### 2.4 Quantifying the impact of parasite nutrient supply on RBC supply and clearance

To examine whether RBC supply and clearance dynamics change with parasite nutrient treatment (and hence parasite growth rate), while accounting for intra- and inter-mouse variation, we performed the following analysis (Figures 2C-D). The data-transformation scheme yielded a distribution of 2000 supply and clearance trajectories for each mouse (Figure 2B), each with an associated likelihood (e.g., Figure 2C, 300 trajectories displayed for ease of interpretation). We calculated the relative likelihood of each trajectory by comparing each likelihood to the maximum likelihood trajectory for a given mouse. Finally, grouping the trajectories by treatment, we used the `wquant` function from the **pomp** package (King et al., 2016) to calculate weighted quantiles (5%, 50% and 95%), using the relative likelihoods as sampling weights (Figure 2D). We thus generated a median trajectory for each treatment that we can use to capture the impact of parasite nutrient supply on the RBC supply and clearance responses, while accounting for uncertainty in the fitting of the data-transformation scheme.

### 2.5 Regression analysis of the RBC supply function

Visualization of the relationship between reticulocyte density at time *t*, *R_t_*, and several lagged values of total RBC density (*RBC_t−i_*, where *i* = 1, 2, 3, 4, 5 and *RBC_t_* = *R_t_* + *E_t_*) suggested that the relationship between *R_t_*and *RBC_t−i_* might be best described by two functions rather than one (cf. §3.1, Figure 3). Specifically, visual analysis suggested that (i) reticulocytes are supplied in at least two distinct phases and (ii) their supply dynamics may change with parasite nutrient treatment. To investigate these possibilities, we built a suite of regression models differing in (a) the form (linear, quadratic, sigmoidal) of the function relating reticulocyte supply *R_t_*and total, lagged RBC density *RBC_t−i_* (*i* = 1, 2, 3, 4 or 5), (b) the phasic nature of said function (i.e., whether the response was mono- or biphasic), (c) the day post-infection on which—in the case of biphasic models—the two phases begin/end (i.e., the breakpoint; either 8, 9, 10 or 11), (d) the effect of parasite nutrient treatment (pABA).

**Figure 3.**
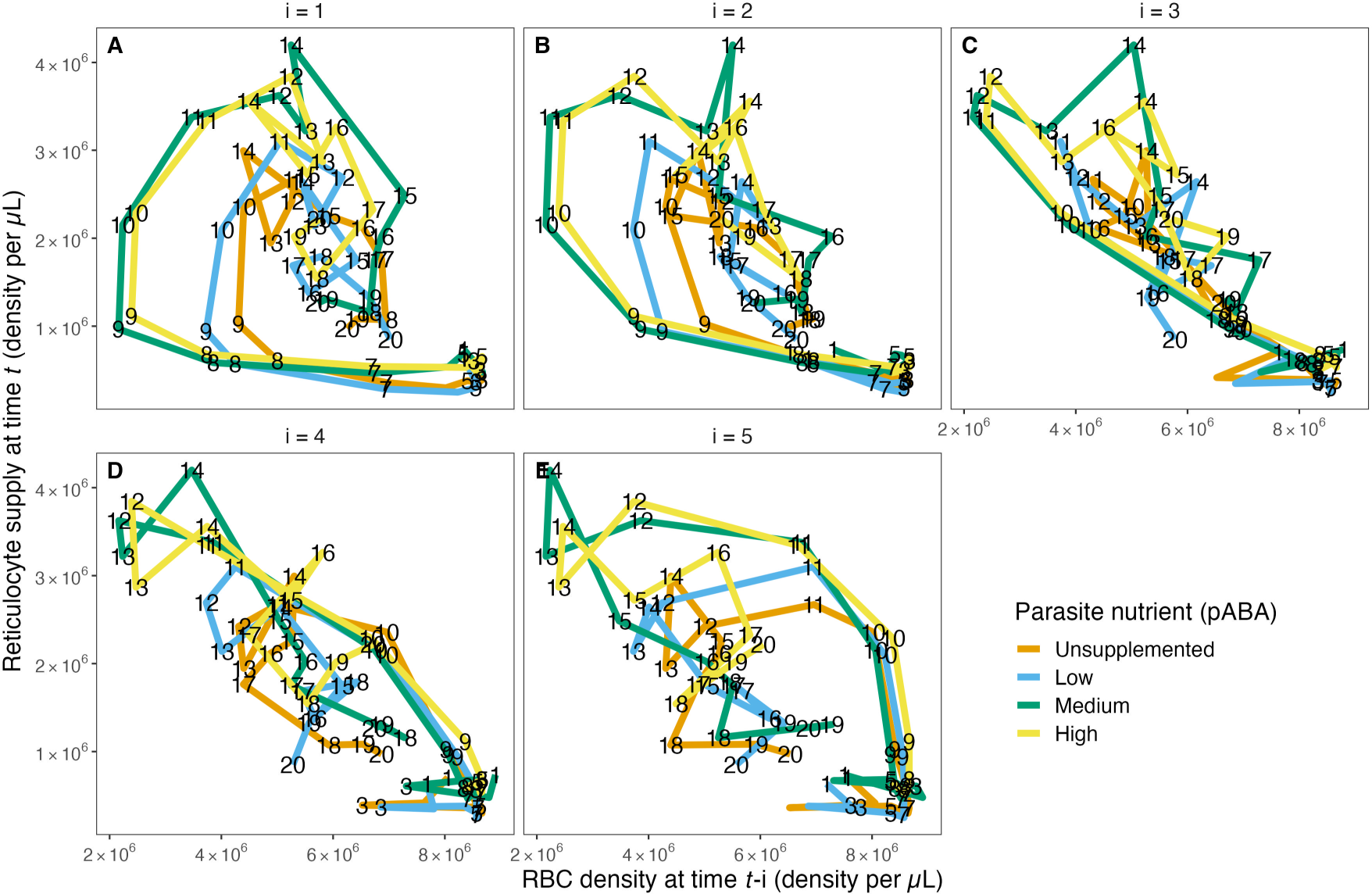
The reticulocyte supply function does not take the form of a single function, as is commonly assumed. The relationship between RBC density on day *t −* 1 (A), *t −* 2 (B), *t −* 3 (C), *t −* 4 (D) and *t –* 5 (E) and reticulocyte density on day *t* for each of the four parasite nutrient (pABA) treatments. The paths are labelled with time *t*, i.e., a point on the path in (A) labelled 8 shows the reticulocyte supply on day 8, given the RBC density on day 7. A point on the path in (B) labelled 8 shows the reticulocyte supply on day 8 given the RBC density on day 6. Panels (C-E) work analogously but for RBC densities lagged by 3, 4 and 5 days.

In total, we compared 160 models. We tested 8 model forms (Table 4): 3 quadratic (Models A-C), 3 linear (Models D-F) and 2 sigmoidal (Models G-H). For the monophasic case, we considered all 8 model forms with 5 lags (*RBC_t−i_* where *i* = 1, 2, 3, 4 or 5). For the biphasic case, we considered Models A-F (i.e., we only considered the case of a monophasic sigmoidal form) with 5 lags and 4 breakpoints.

**Table 4.**
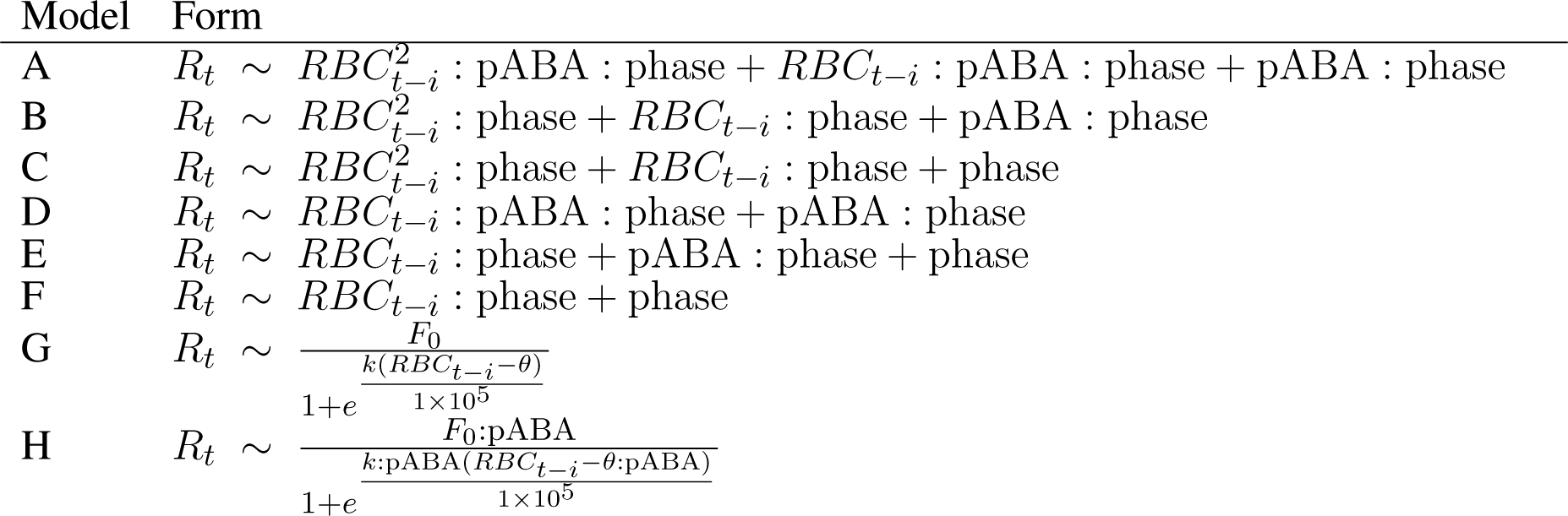
Model forms considered for the relationship between *R_t_* and *E_t−i_*, where *i* = 1 − 5. For each of the five time lags (*i* values) considered, we fit 32 models: 8 monophasic models of forms A-H plus biphasic versions of models A-F with 4 different breakpoints (i.e., days 8, 9, 10 or 11 post-infection; 6 models × 4 breakpoints = 24 models). Note that when no breakpoint is specified, the phase term becomes irrelevant so that, for example, Model F becomes equivalent to the monophasic, linear model commonly used in previously published predictive models (cf. Figure 1). Note that pABA is treated as a discrete factor. Also note that to address the issue of multicollinearity associated with regressing against higher order polynomials, we opted to use orthogonal polynomials for fitting model forms A-F. We considered two different sigmoidal models of similar form: Model G assumes the maximum reticulocyte supply *F*_0_, the RBC density where reticulocyte production is half its maximum *θ* and the steepness coefficient *k* do not vary with pABA treatment, while Model H assumes each pABA treatment has a unique value for these three parameters.

| Model | Form |
| --- | --- |
| A | $R_t \sim RBC_{t-i}^2 : \text{pABA} : \text{phase} + RBC_{t-i} : \text{pABA} : \text{phase} + \text{pABA} : \text{phase}$ |
| B | $R_t \sim RBC_{t-i}^2 : \text{phase} + RBC_{t-i} : \text{phase} + \text{pABA} : \text{phase}$ |
| C | $R_t \sim RBC_{t-i}^2 : \text{phase} + RBC_{t-i} : \text{phase} + \text{phase}$ |
| D | $R_t \sim RBC_{t-i} : \text{pABA} : \text{phase} + \text{pABA} : \text{phase}$ |
| E | $R_t \sim RBC_{t-i} : \text{phase} + \text{pABA} : \text{phase} + \text{phase}$ |
| F | $R_t \sim RBC_{t-i} : \text{phase} + \text{phase}$ |
| G | $R_t \sim \frac{F_0}{1 + e^{\frac{k(RBC_{t-i} - \theta)}{1 \times 10^5}}}$ |
| H | $R_t \sim \frac{F_0 : \text{pABA}}{1 + e^{\frac{k : \text{pABA} (RBC_{t-i} - \theta : \text{pABA})}{1 \times 10^5}}}$ |

We fit these statistical models to the reticulocyte supply and total RBC trajectories, as estimated from the data-transformation scheme. To account for variation in the shape of the 2000 trajectories outputted in the scheme-fitting process (Figures 2B, S3, *Supplementary Material*), we used a bootstrapping approach. Specifically, we fit each of the 160 regression models to 1000 “datasets”, each of which contained a single trajectory from each of the ten mice. To construct each dataset, we sampled one trajectory per mouse such that the probability that a trajectory was sampled was proportionate to its relative likelihood. The models were then fit to each dataset in turn and the best fit model selected via AICc. Note that to address the issue of multicollinearity associated with regressing against higher order polynomials, we opted to use orthogonal polynomials for model fitting.

### 2.6 The dynamics of RBC clearance rate and reticulocyte supply during infection

To investigate whether (i) clearance rate of uninfected RBCs 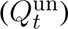 and (ii) reticulocyte supply changes through time and/or with parasite nutrient (pABA) treatment, we again performed a bootstrap analysis. For the analysis of clearance rate dynamics, we compared five generalized additive models (`gam`) that described *Q*^un^ as: (A) a constant function; (B) a smooth function of time, i.e., day post-infection; (C) a smooth function of time and pABA treatment; (D) a linear function of RBC density (per the commonly posited models); or (E) a linear function of RBC density and pABA treatment (Table 5). For the analysis of reticulocyte dynamics, we fit only the first three models (Table S1, *Supplementary Material*). Again, the models were fit to 1000 datasets, each composed of ten trajectories and sampled as described above. Model fitting was performed using the **mgcv** package in **R** (Wood, 2017; R Core Team, 2022); model selection was performed using AIC.

**Table 5.**
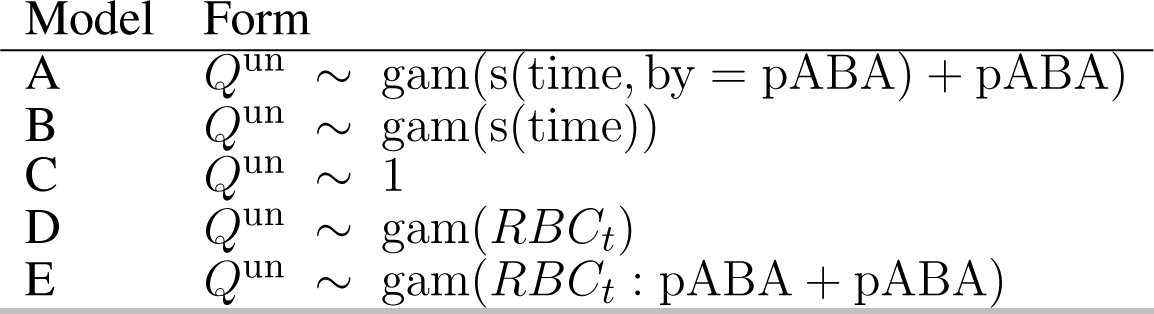
Model forms considered for predicting clearance rate *Q*^un^ as a function of time and a parasite nutrient pABA. Model fitting was performed using the `gam` function from the **mcgv** package (Wood, 2017) in **R** (R Core Team, 2022), which fits a generalized additive model to data.

## 3 RESULTS

Host supply and destruction of uninfected RBCs are major drivers of the availability of RBCs during infection. We set out to examine (i) whether mathematical functions that are used to describe these processes (Tables 1, 2; Figure 1) capture our data and (ii) whether they change with an experimental treatment (dietary supply of a parasite nutrient) that impacts parasite growth rate and dynamics (Figures S1, S2), but has no effect on the host (Fenton et al., 1950). To do so, we analyzed time-series data using a data-transformation scheme that allows us to quantify host RBC supply and clearance responses through time. Specifically, this analysis generated a distribution of response trajectories for each mouse, each of which was associated with a likelihood (Figure 2B). By analyzing this distribution we can quantify the shape of the functions that describe reticulocyte supply and uninfected RBC clearance while accounting for the fact that multiple realizations of the data-transformation scheme may explain the data.

### 3.1 The reticulocyte supply response is not well described by a monophasic function

Reticulocyte (i.e., newly supplied RBC) supply is often modeled as a single, decreasing function of (lagged) RBC density or as a constant (Figure 1A). These models hypothesize that, when plotted, the relationship between reticulocyte density, *R_t_*, versus (lagged) RBC density will take the shape of a single curve.

Contrary to this hypothesis, we found that—broadly—the shape of the *R_t_*-*RBC_t−i_* curve takes the form of a loop (Figure 3A-B,E), suggesting that reticulocyte supply may not be well described by a single function of any (tested) form. We found the curves relating *R_t_* to *RBC_t−_*_1_, *RBC_t−_*_2_ and *RBC_t−_*_5_ take a looped form. Interestingly, however, the shape of the relationship between *R_t_* versus *RBC_t−_*_3_ and *R_t_*versus *RBC_t−_*_4_ curves remained unclear upon initial visual inspection. Plots of the raw data (Figure S8, *Supplementary Material*), rather than the output of the data-transformation scheme, showed the same patterns. Together, these graphical analyses suggested that reticulocyte supply may not occur according to a single, monophasic function of RBC density but rather may be better described by two distinct phases that start/end roughly halfway through the first 20 days post-infection.

To test this hypothesis, we fit a suite of 160 models to each of 1000 iterations of our data-transformation scheme. Together, these models allowed us to ask: (i) whether reticulocyte supply is better described as biphasic or monophasic, (ii) if it is biphasic, what time (i.e., day post-infection) represents the best “breakpoint” between the two phases, (iii) whether *R_t_* supply is better described by a linear, curved or sigmoidal function of *RBC_t−i_* (where *i* = 1 − 5) and (iv) whether *R_t_* supply changes with supplementation of a nutrient (pABA) that changes parasite growth rate (Table 4).

Across the 160 models considered, we found overwhelming evidence that reticulocyte supply is not well described by a monophasic function and is better described using two phases. Out of 1000 iterations of our regression analysis, a monophasic model was selected only 3.2% of the time (Figures 4A, S4, *Supplementary Material*). Instead, our analysis instead implies that reticulocyte supply is better described in two phases that begin/end at day 10 post-infection (breakpoint selected 54.0% of the time). That said, day 9 (23.7%) and day 11 (14.3%) were supported a substantial portion of the time.

**Figure 4.**
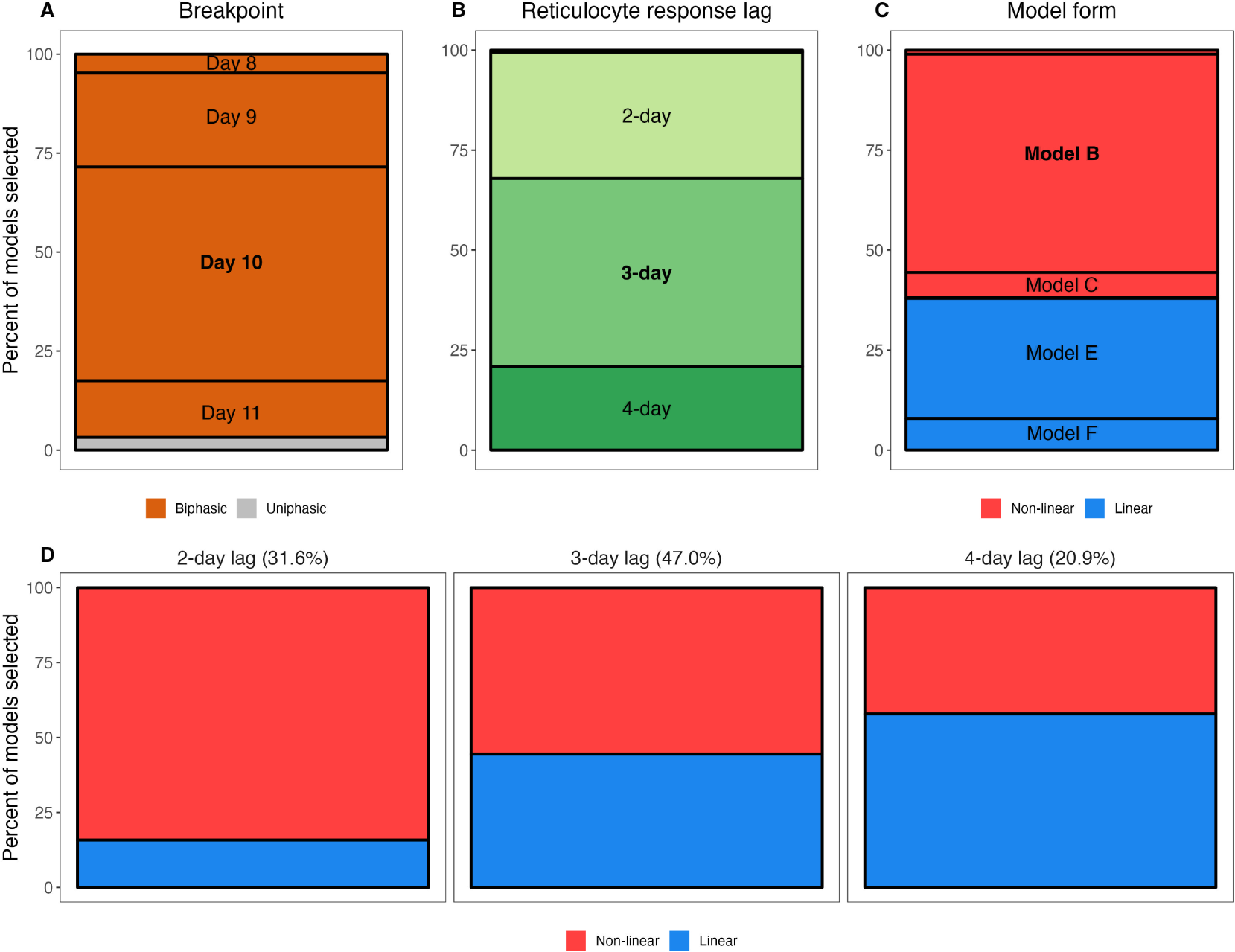
Qualitative features of the statistical models that were selected in our bootstrapped regression analysis of the RBC supply response. A-C) Marginal percentages of breakpoint (A), reticulocyte response lag (B) and model form (C). For example, (A) shows that out of 1000 selected models from the regression analysis, 4% had a breakpoint at day 10 post-infection. Note that in (B), a 5-day lag is not visible as it was never selected and a 1-day lag accounted for only 0.5% of models selected. Similarly, in (C), Models A and D are difficult to visualize as they account for 1.0% and 0.2% of models selected, respectively, while Models G and H (sigmoidal model forms) are not included as they were never selected. D) Marginal percentages of general model form (non-linear versus linear) for models specifying a 2-day, 3-day or 4-day lag. Percentages at the top of each panel reflect the percent of models selected with a 2-day lag, 3-day lag and 4-day lag.

Model comparison did not indicate strong support for a particular lag or model form, however. Our analysis suggests that 1- and 5-day response lags do not describe the data well (i.e., they were selected 0.5% and 0% of the time, respectively), but models with 2, 3- or 4-day lags were all selected at high frequencies (31.6%, 47%, 20.9%, respectively; Figure 4B). Similarly, we find weak support for a non-linear reticulocyte response: 61.9% of the selected models were non-linear (Models A-C; Table 4; Figure 4C). Notably, and in accordance with the observation that the linearity of the reticulocyte supply “loop” increases with the response-lag (Figure 3), we found a systematic relationship between the lag and linearity parameters of the selected models (Figure 4D). For example, of the models that were selected as the “best” description of the data and which specified a 2-day lag, 84.2% were non-linear. However, the frequency with which non-linear models were selected declined with increasing lag, halving to 42.1% in the case of models with 4-day lags (Figure 4D)

### 3.2 Parasite nutrient supply changes the magnitude but not the shape of the reticulocyte supply response

Our analysis strongly suggests that dietary supply of parasite nutrients (pABA) alters the baseline concentration of reticulocytes supplied during each phase of the infection (Models B and E selected in 84.6% of the iterations). Parasite nutrient supply does not appear to change the rate at which reticulocyte supply increases as RBC density falls: models that posit an effect of pABA on the slope/shape of the response (A, D) were selected just 1.2% of the time.

The effect of parasite nutrient (pABA) supply is more pronounced in the second phase of the infection than the first, according to the majority of selected models (Figures 5, S5). For example, per our most-frequently selected model, in the first phase of infection, unsupplemented mice with an RBC density of 5 × 10^6^ supply the bloodstream with 1.58 × 10^6^ reticulocytes. Mice supplemented with the highest concentration of pABA, and experiencing the same degree of anemia, supply the blood with 1.73 × 10^6^reticulocytes, i.e., 11% more than unsupplemented mice. In the second phase, the difference among mice in different nutrient treatments is more marked: mice in the high pABA treatment with an RBC density of 5 × 10^6^ resupply the blood with 28% more reticulocytes than those in the unsupplemented pABA treatment. Notably, all but two of the nine most-frequently selected models (which together comprise 76.2% of all models selected) posit that parasite nutrient supplementation alters RBC supply in this way (Figure S5). The fifth and sixth most-frequently selected models also suggest that there is a greater effect of pABA in the second phase but differ from the other models among the top nine by suggesting reticulocyte supply is elevated in the first phase relative to the second for the unsupplemented and low pABA treatments, while reticulocyte supply under the medium and high pABA treatments do not differ greatly between phases (Figure S5E-F). This difference is likely because these models posit a late breakpoint (day 11) and long response-lag (4 days). In conclusion, our results broadly suggest that for each RBC that is destroyed, mice supplemented with the two highest concentrations of pABA add more reticulocytes to the bloodstream than those supplemented with the two lower concentrations and this effect is most pronounced in the second phase of infection.

**Figure 5.**
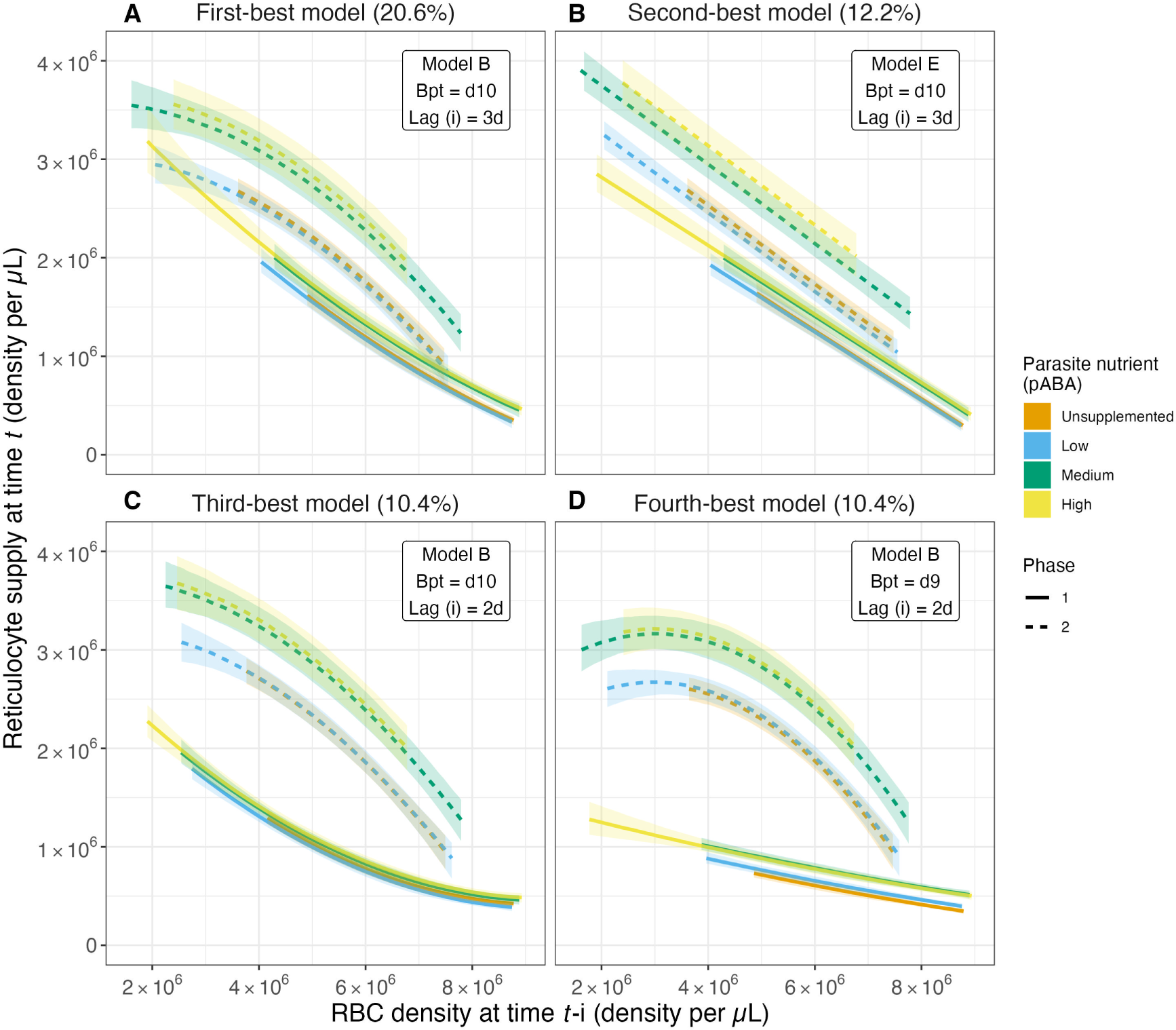
The magnitude of the reticulocyte supply response changes with parasite nutrient supply. Predicted values of reticulocyte response as a function of lagged RBC (red blood cell) density for the top four models from the reticulocyte response regression analysis. These top four models account for 53.6% of all models selected. Solid and dashed series reflect median values of the predicted reticulocyte response obtained from calculating predicted values for a given model from each of the 1000 regression analysis iterations. Ribbons along solid and dashed lines provide the 80% confidence intervals. Colors reflect the parasite nutrient (pABA) treatment. Solid versus dashed series reflect median values in the first versus second phase of acute infection, where the days at or below the breakpoint are considered the first phase and those after the breakpoint are considered the second phase. Percentages in the titles of (A-D) reflect the percent of all models selected that are of the specified form. For example, 20.6% of all models selected followed model form Model B (Table 4), had a breakpoint at day 10 post-infection and specified a reticulocyte response lag of 3 days.

For the sake of completeness, we also fit our 160 models to the raw reticulocyte and RBC data. The best-fitting model in this analysis was the same as the second most-frequently selected model in the analysis of model trajectories. It suggests that : (i) reticulocyte response is better described by a biphasic function rather than a monophasic function, with each phase beginning/ending on day 10, (ii) reticulocyte supply is better described by a linear function (rather than a quadratic or sigmoidal function) of RBC density 3 days earlier and (iii) parasite nutrient supply alters the magnitude of reticulocyte supply, particularly in the second phase of the infection (Figure S9).

### 3.3 Reticulocyte supply varies with time and parasite nutrient supply

To further investigate the impact of the parasite nutrient pABA on reticulocyte supply dynamics during infection, we compared a suite of generalized additive models (Table S1, *Supplementary Material*). This analysis strongly suggests that reticulocyte supply varies with both time and pABA treatment (Model A, winning best support in 95.9% of model selection iterations). Specifically, reticulocyte concentrations are higher prior to day 7, and reach a higher peak, in mice supplemented with pABA (i.e., with fast growing parasites; Figures S10A-B)

### 3.4 RBC clearance rate is not well described by commonly employed functions

As with RBC supply, we found that commonly employed functions (Figure 1B-C) poorly describe the dynamics of RBC clearance during infection. Visualization of the RBC clearance trajectories suggest that RBC clearance varies markedly with time, beginning at a relatively constant rate, before increasing sharply and peaking around day 11-12 post-infection. In addition, the graphical analysis suggested that clearance rate varies with the supply of parasite nutrient (Figure 6A). To investigate this hypothesis, we again performed a bootstrapped regression analysis. We found that the model forms reflecting common assumptions of RBC clearance (Models C-E, Table 5) were not supported and indeed never appeared as the best model form in our regression analysis (Figure 6B). Rather, we found the remaining two models (Models A & B, Table 5)—which specified clearance as a function of time—outperformed the aforementioned models at roughly equal frequency (Model A at 52.3%, Model B at 47.7% frequency). According to the best-supported model (Model A), the zenith in clearance rate is higher in mice supplemented with high concentrations of parasite nutrients (Figure 6A). These results suggest that clearance rate is not well described as a constant or as a linear function of RBC density.

**Figure 6.**
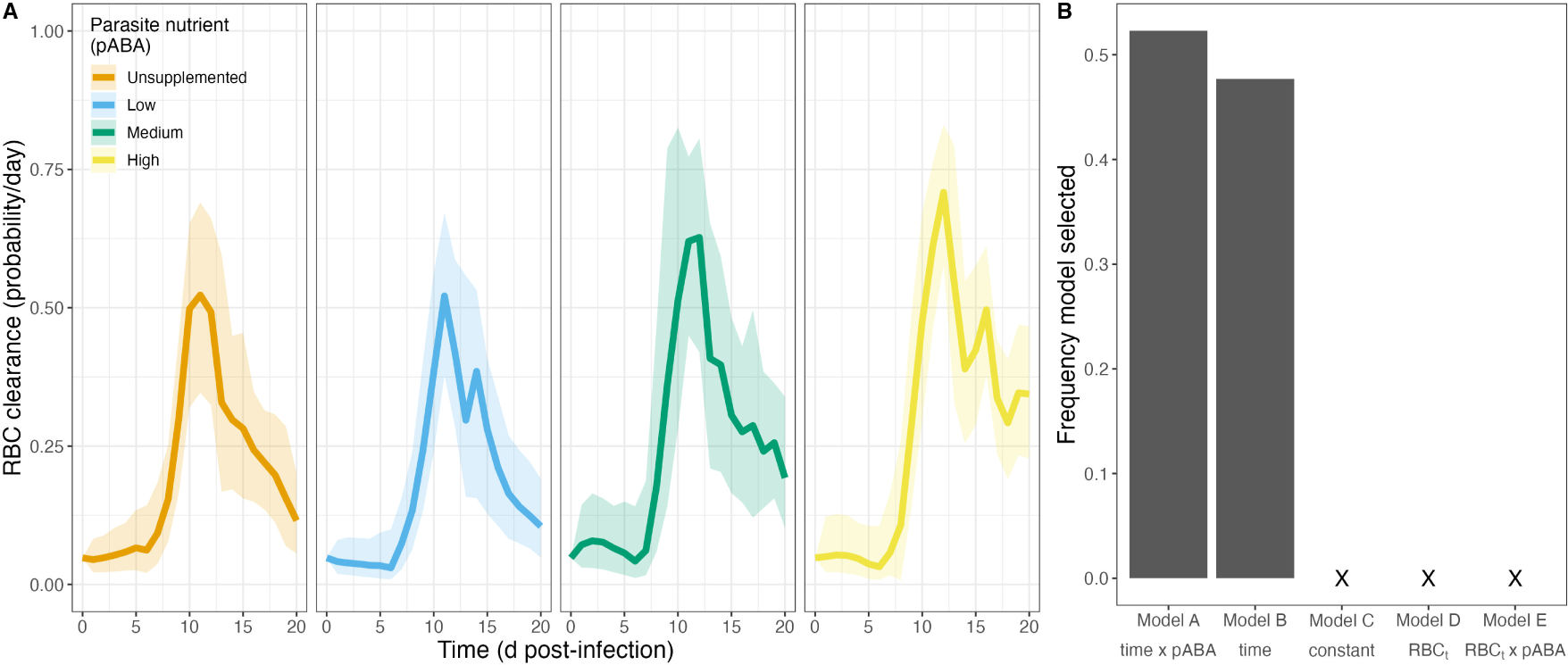
Clearance rates of uninfected RBCs during *P. chabaudi* infection vary through time and with a parasite nutrient. A) Median treatment-level red blood cell (RBC) clearance rates over time, in each of four groups of mice that received a different concentration of the parasite nutrient pABA. B) Frequency of 5 models selected via AIC. Note that a model including a linear relationship between clearance rate and *RBC_t_* was never selected.

## 4 DISCUSSION

Mathematical models have shaped our understanding of diverse phenomena in malaria biology—from the dynamics of infections (Haydon et al., 2003; Wale et al., 2019; Kochin et al., 2010; Mideo et al., 2008), to the evolution of transmission investment (i.e., gametocyte production) (Greischar et al., 2016) and the design of control interventions (Camponovo et al., 2021). In this work, our goal was to interrogate commonly held assumptions about the functional forms of red blood cell (RBC) supply and clearance. We emphasize that our intent was not to propose the “correct” functional forms of RBC supply and clearance function and acknowledge that some recent studies have used functions other than those investigated here (Tables 1, 2). Here, we explicitly were interested in whether the aforementioned commonly held assumptions are consistent with data on how both mature and immature RBC densities change with time and with parasite growth rate, as manipulated by parasite nutrient supply. We found that common model assumptions about the processes that regulate RBC abundance during infection do not hold. First, we found that RBC supply is not well described as a single-valued function of RBC deficit as is commonly assumed (Figure 1A, Table 1). Rather, it is better described as two functions that describe roughly the two halves of the acute phase of infection (Figure 4A-B). Second, and similarly, we found that RBC clearance is poorly described by constant or linear functions (Figure 6B), as has frequently been postulated (Figure 1B-C, Table 2). Rather, models specifying that RBC clearance rate changes with time perform better (Figure 6). This result suggests that either clearance rate explicitly depends on time *per se* or, more likely, that it is a function of one or more unmeasured entities that vary with time. Finally, our analysis suggests that RBC supply and clearance may change with parasite growth rate (as manipulated by pABA; Figures 5, 6, S10). pABA is a parasite nutrient, as it i) increases parasite growth rate (Figure S2), but is not a nutrient for the host (Fenton et al., 1950). Accordingly, the effect of pABA on host RBC production is likely to be indirect i.e., by increasing parasite growth rate it stimulates higher erythropoiesis. How this phenomenon comes about is unclear and warrants investigation.

Our finding that RBC (i.e., reticulocyte) supply is better described by two, differently shaped functions (rather than one single function) is consistent with recent immunohematological studies of reticulocytosis. These studies provide us with testable, mechanistic explanations for our findings vis-a-vis reticulocyte supply. It is now widely recognized that the inflammatory response, as stimulated by infection, causes “stress erythropoiesis” (reviewed in Ruan and Paulson (2023)). This response is characterized by two phases: first, there occurs a cytokine-mediated suppression of erythropoiesis and an acceleration of erythrophagocytosis; second, there occurs a “pulse” of reticulocyte production in the spleen, which is stimulated by a heme-mediated signaling cascade (Yap and Stevenson, 1992; Chang and Stevenson, 2004; Paulson et al., 2020; Bennett et al., 2019). Given (i) that malaria infection induces significant inflammation, erythrophagocytosis and heme-accumulation in the spleen and (ii) the concordance between the timescale of stress erythropoiesis and our RBC supply (i.e., reticulocyte supply) curve (Figure S10), it is reasonable to hypothesize that similar mechanisms underlie the biphasic reticulocyte supply dynamics we have observed.

Stress erythropoiesis may also drive the increase in RBC clearance in the post-peak phase of infection. In addition to stimulating an increase in the production of reticulocytes, stress erythropoeisis is associated with an increase in the turnover of RBCs (Libregts et al., 2011; Klei et al., 2019; Fernandez-Arias et al., 2016; Mourão et al., 2018). Other, non-mutually-exclusive mechanisms may also be at play. Alternatively, infection may stimulate an increase in “normal” senescence-mediated RBC clearance by changing the process of RBC aging. For example, inflammation-associated oxidative stress may increase the speed at which RBCs senesce or—as in birds infected with malaria—reticulocytes produced during infection may enter the bloodstream “biologically older” than those produced under conditions of health (Asghar et al., 2015). Unfortunately, because of the difficulties in aging mammalian RBCs (which, unlike bird RBCs, do not possess telomeres), the latter hypothesis is at present difficult to test.

Our analysis also implies that parasite nutrient (pABA) supplementation (and by extension, pathogen growth rate) changes the “rules” by which the RBC population is regulated. This novel finding may explain previous observations vis-a-vis the impact of pABA supplementation on the dynamics and disease of malaria infections. We (and others) have observed that infections of mice supplemented with pABA reach higher peak densities (or parasitemias) and are more likely to exhibit a second wave of growth in the post-peak phase of infection (Hawking, 1954; Jacobs, 1964; Morgan, 1972; Wale et al., 2017b,a). Hitherto, our explanation for this observation has been that pABA changes the growth rate of parasites, likely by changing the number of merozoites each parasite produces (i.e., “burst size” (Tan-Ariya and Brockelman, 1983)). Our results imply that pABA-supplementation promotes parasite growth via a second mechanism, by increasing the amount of resource that can be exploited, i.e., the “carrying capacity” of the bloodstream. In addition, we have also shown that pABA-supplementation alters the “tolerance” of mice in the second phase of infection (from c. 10 dpi) (Wale et al., 2017b). Specifically, supplemented mice have more RBCs in their bloodstream than unsupplemented mice at the same parasite burden. This phenomenon may also be explained by the increase in magnitude reticulocyte supply, which we found to be particularly profound during the second phase of the infection (although it may be outweighed by the concomitant increase in RBC clearance). Further analysis should be conducted to (i) verify our finding that RBC supply (i.e., erythropoiesis) and RBC clearance change with parasite nutrient supply (given the small size of the experiment conducted here) and, (ii) quantify the net effect on parasite population dynamics and the traits that mediate them. In particular, we anticipate that parasite nutrient supply could impact the dynamics of asexual parasites (via its impact on burst size) and sexual-stage parasites (i.e., gametocytes). Both the rate at which malaria parasites commit to gametocyte production (i.e., conversion rate) and the sex ratio of gametocytes varies with reticulocytosis (Gautret et al., 1996; Reece et al., 2005; Birget et al., 2017). Hence, if parasite nutrient supply indirectly impacts reticulocytosis, as our results suggest, it could contribute to plasticity in malaria parasites’ conversion rate (Birget et al., 2019).

The conclusions we make from mathematical models of malaria infection dynamics and evolution could change if, as our analysis suggests, RBCs are regulated in a manner not captured by said models. For example, there is a longstanding debate about whether parasite population growth in the blood is primarily controlled by the availability of RBCs (“bottom-up” forces) or by immune-mediated killing (“top-down” forces) (Antia et al., 2008; Kochin et al., 2010; Wale et al., 2019; Haydon et al., 2003). Our analysis implies that RBCs may be more limiting than is often assumed, since most models do not capture the suppression of erythropoiesis at the infection’s outset (i.e., in the first “phase”). Evolutionary inferences may also be impacted by deviations between the way RBC supply is modeled versus how it actually occurs. Evolutionary theory states that—all else being equal—competition for resources (RBCs) impacts selection for virulence-related traits, such as growth rate or the number of merozoites produced per parasite (or burst size) (Frank, 1996; Mosquera and Adler, 1998; Bremermann and Pickering, 1983; van Baalen and Sabelis, 1995; Nowak and May, 1994). Our results imply that not only is RBC limitation potentially stronger than assumed but that all else is not in fact equal, i.e., RBC dynamics systematically vary with parasite traits (growth rate). It is difficult to intuit how such feedbacks will impact within-host competition or virulence evolution but, given the profound evolutionary and health consequences of parasite virulence, their explication is warranted (Alizon and Michalakis, 2015).

We recognize that our inferences, although data-driven, are to some degree shaped by the assumptions of the data-transformation scheme used to generate them. However, relaxation of these assumptions is highly unlikely to negate our central conclusion that RBC dynamics do not behave as commonly modeled. Among the several simplifying assumptions we make, two are most likely to have significant impact on our conclusions. The first is that all RBCs are equally invadable by *Plasmodium chabaudi* (Jarra and Brown, 1989; Yap and Stevenson, 1994; Carter and Walliker, 1975; Taylor-Robinson and Phillips, 1994). If parasites in fact exhibit a preference for a narrower subset of RBCs (e.g., invade only mature RBCs), we will have underestimated parasite-mediated destruction of RBCs (i.e., via *M* or *W*) and, as a result, overestimated the clearance of uninfected cells, *Q*^un^. The dynamics of *Q*^un^ are unlikely to qualitatively change in this event, however. The second assumption that could affect our inference is that reticulocytes mature into erythrocytes after only one day in circulation (Ney, 2011; Gronowicz et al., 1984; Koury et al., 2005; Wiczling and Krzyzanski, 2008; Noble et al., 1989). If reticulocytes remain immature RBCs for longer than 24 hours once in circulation (as some studies imply, e.g., Thakre et al., 2018; Ganzoni et al., 1969; Gronowicz et al., 1984), we will have “double-counted” reticulocytes and hence overestimated RBC supply at each time-point. Nonetheless, we stress that our goal here was not to present a “better”, or even predictive, model of RBC dynamics, but rather to ask whether or not RBC supply and clearance conform to commonly-invoked assumptions. It seems highly unlikely that the many degrees of freedom that would be introduced into models by the addition of more realism would combine to rescue the simplistic assumptions we interrogate here. After all, these assumptions have been made for theoretical convenience and parsimony.

In this paper, we have established a negative result: simplifying assumptions made about RBC dynamics during malaria infections are not consistent with our data. Indeed—and interestingly—our findings suggest that RBC supply cannot be well-described as a single-valued function of RBC concentration. The implication is that new mathematical models are needed to fully explain and predict the dynamics of malaria infections. How might we achieve this goal?

We propose two directions for future research. First, over the short-term, richer time-series datasets could help us to quantify at-present poorly identified model parameters or processes. Our analysis exemplifies the value of such efforts. Historically, it has not been possible to disentangle whether a host is accelerating the clearance of RBCs (“bystander killing”) or simply not supplying the bloodstream with RBCs: both processes result in a deficit of RBCs (as discussed in, e.g., Metcalf et al., 2011, 2012). Simply by measuring reticulocyte dynamics, we were able to disentangle these processes. We thus advocate for experimental efforts to measure the various parameters that define the RBC supply and clearance functions but whose values are at present not well estimated. For example, both the the maturation rate of reticulocytes and the clearance rate of RBCs could be measured via the “double-labeling” of RBCs (as in Gifford et al., 2006; Klei et al., 2019; Bennett et al., 2019). Experiments in bird hosts, whose RBCs come “prelabelled” with telomeres, could be similarly informative. It is likely that with more informative datasets we could achieve our second, long-term goal: to replace explicitly time-dependent models of the mechanisms that drive RBC dynamics (Table 1, 2) with truly dynamical models. Ultimately, dynamical models will enable us to explain and predict the population dynamics of malaria parasites and the red blood cells they infect.

## AUTHOR CONTRIBUTIONS

All authors contributed to the conceptualization of the manuscript, design of the methodology and design of the visualization. MAEP wrote the manuscript with AAK and NW contributing substantial revision. MAEP and AAK performed the analyses, and MAEP performed the visualization of the methods and results.

## ACKNOWLEDGMENTS

We are thankful to Andrew Read and his laboratory at Penn State University, for performing the experiment that generated data underlying the results reported here and by Wale et al. (2019).

## SUPPLEMENTAL DATA

Supplementary information, figures and tables are available in the *Supplementary Material* document.

## DATA AVAILABILITY STATEMENT

The datasets as well as all code for analyses and figures for this study can be found on GitHub at https://github.com/kingaa/malaria-rbc-dynamics. An archival version of these codes will be stored on Zotero upon acceptance of this paper for publication.

