## Supplementary Material for "Red blood cell dynamics during malaria infection challenge the assumptions of mathematical models of infection dynamics"

### 1 SUPPLEMENTARY TABLES AND FIGURES

#### 1.1 Tables

**Table S1. Model forms considered for predicting reticulocyte supply, $R$, as a function of time and pABA.** Model fitting was performed using the *mcgv* package (Wood, 2017) in **R** (R Core Team, 2022).

| Model | Form |
|---|---|
| A | R∼gam(s(time, by = pABA)+pABA) |
| B | R∼gam(s(time)) |
| C | R∼1 |

#### 1.2 Figures

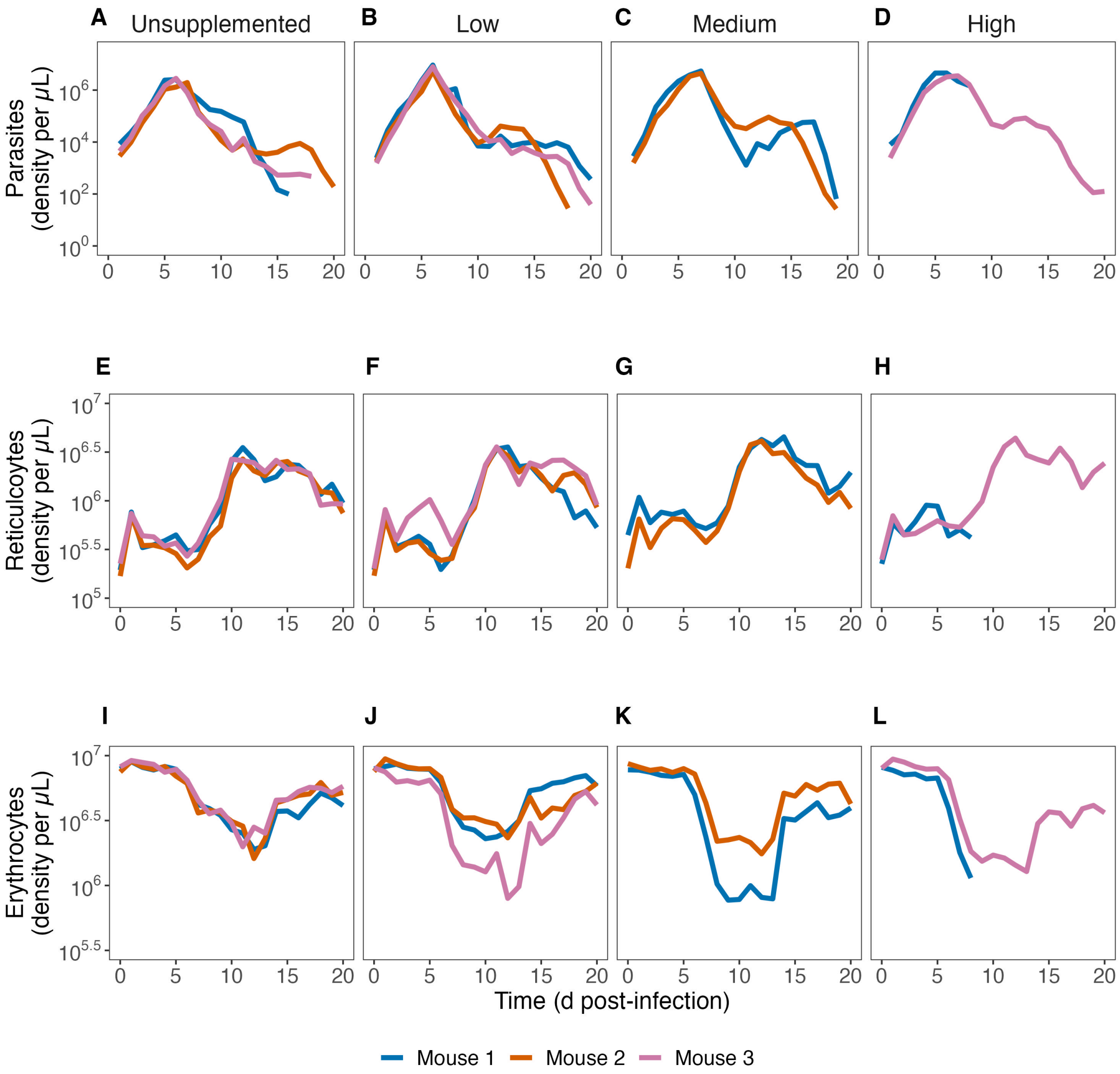


**Figure S1. Raw parasite and RBC density data.** Parasite, reticulocyte and erythrocyte densities for all mice over the first 20 days of infection. Columns reflect pABA treatment (i.e., mice in the unsupplemented treatment received $0\%$ pABA, those in the low treatment received $0.0005\%$, those in the medium treatment received $0.005\%$ and those in the high treatment received $0.05\%$).

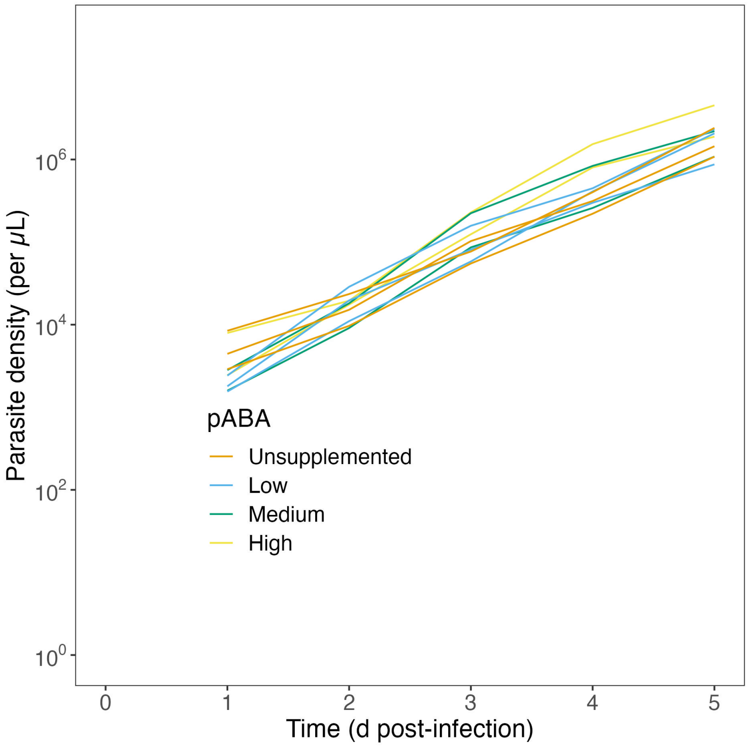


**Figure S2. Density per microliter of parasites over the first 5 days post-infection.** Each line indicates the dynamics in an individual mouse. Mice in the unsupplemented treatment (orange) received $0\%$ pABA, those in the low treatment (blue) received $0.0005\%$, those in the medium treatment (green) received $0.005\%$ and those in the high treatment (yellow) received $0.05\%$

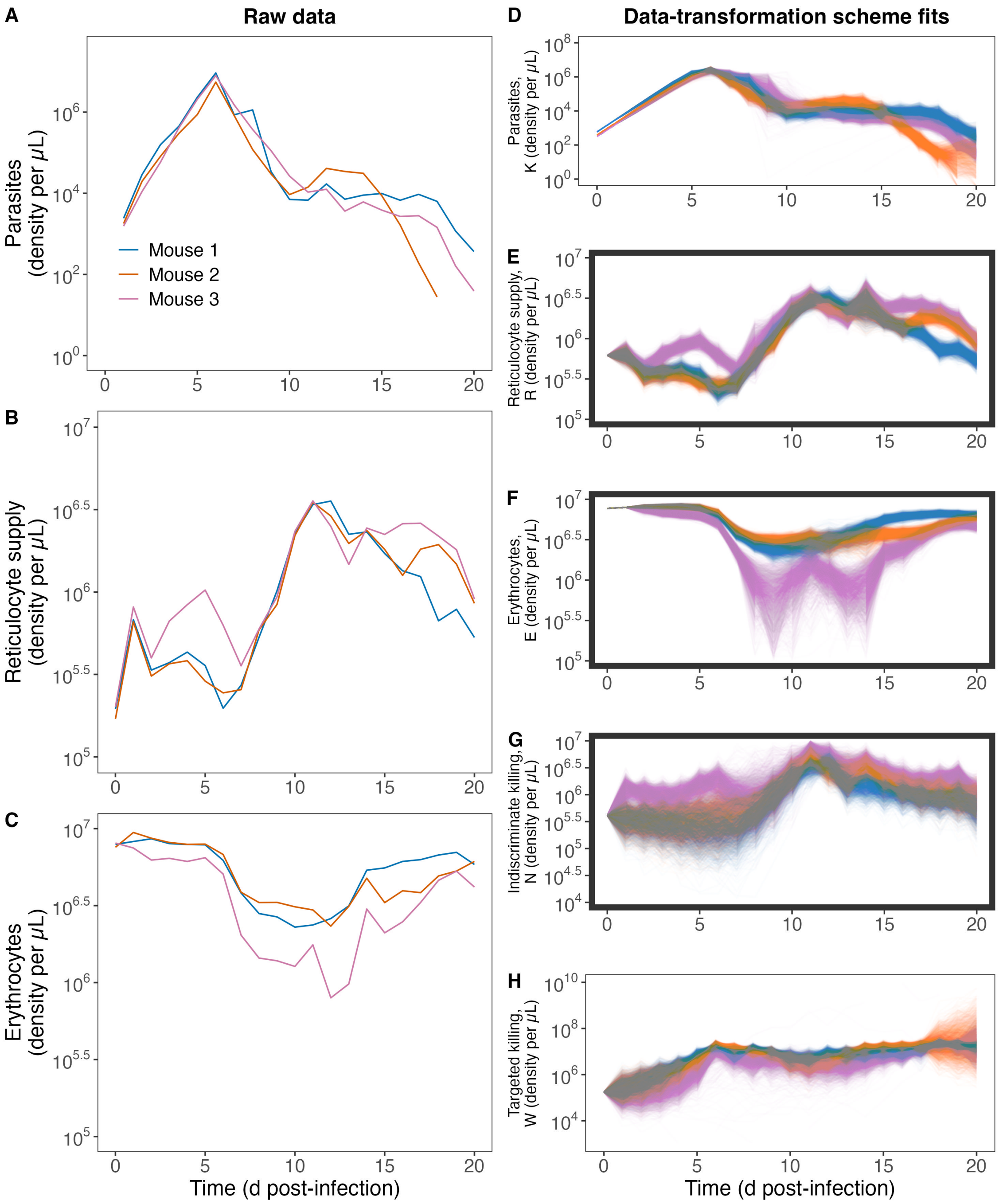


**Figure S3. Example raw data and data-transformation scheme fits for three mice.** (A–C) Parasite, reticulocyte and erythrocyte densities for three mice from the “low” pABA treatment over the first 20 days post-infection. (D–H) Data-transformation scheme trajectories for the same three mice as shown in (A–C). Note, for our analyses we use the trajectories of reticulocyte supply (E), erythrocyte density (F) and indiscriminate killing (G) (bolded panels).

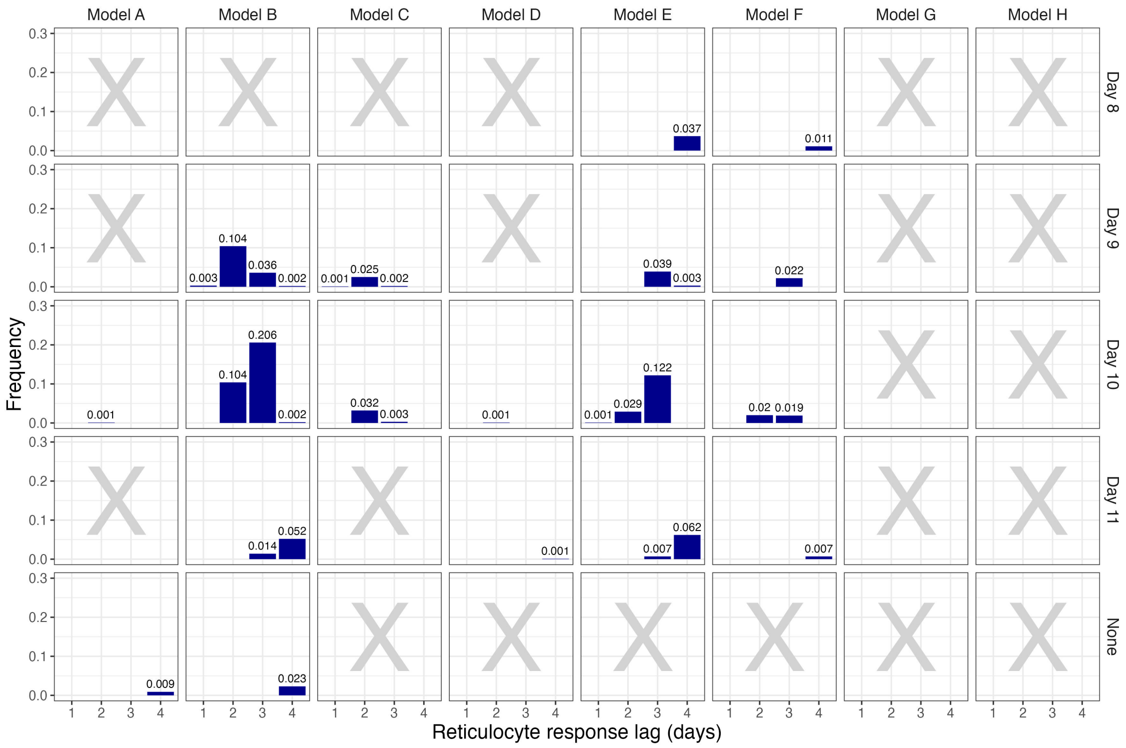


**Figure S4. Frequency with which each model of reticulocyte supply was selected in 1000 iterations.** Rows indicate the breakpoint specified, columns the form of the model (see Table 4, *Main text*). Reticulocyte response lag (in days) is displayed on the x-axis of each panel. The 5-day lag has been omitted as models that specify this lag were never selected. A gray cross in a given panel indicates that none of the models with this combination of model form (columns) and breakpoint (rows) was ever selected.

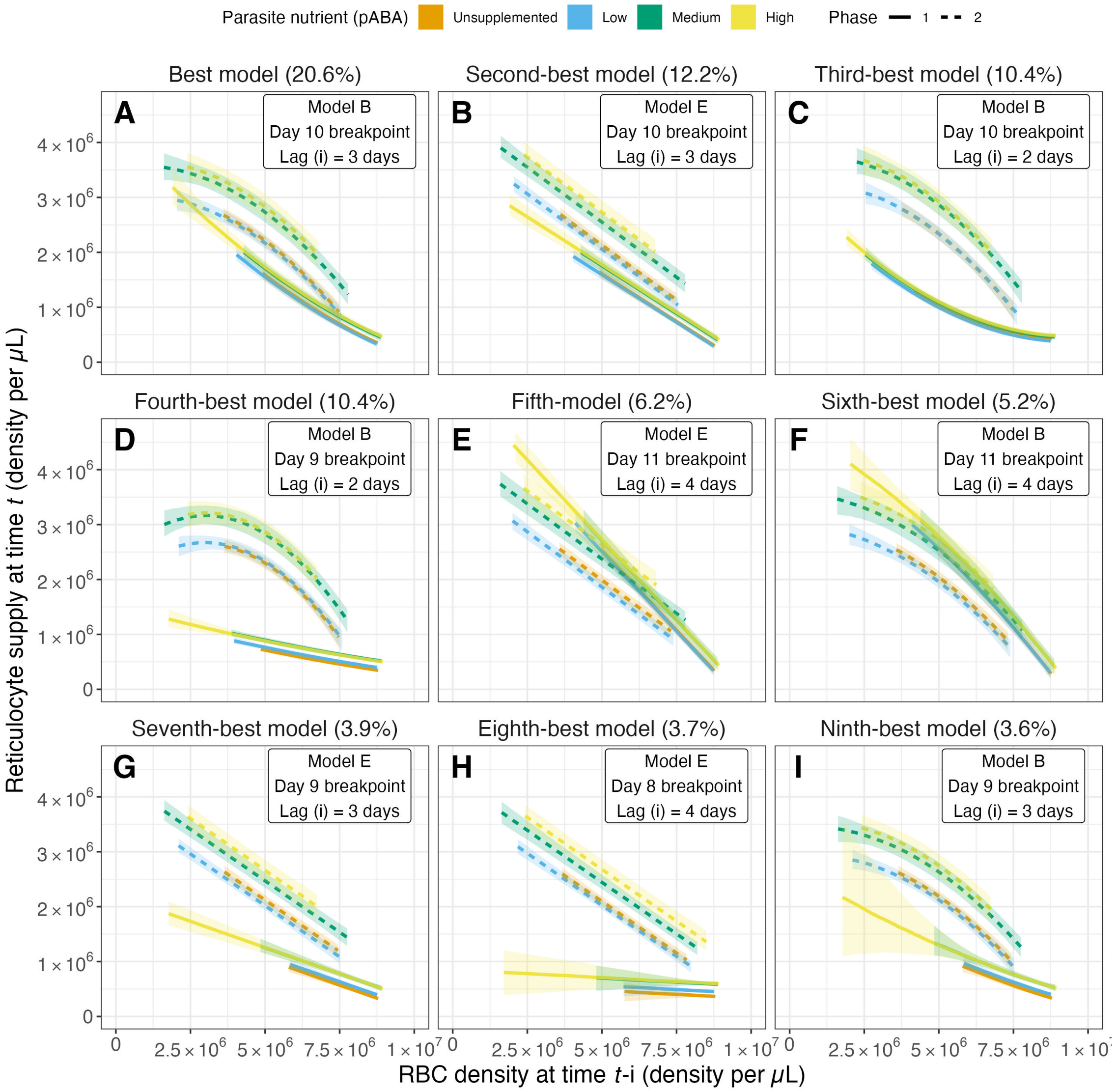


**Figure S5. The reticulocyte supply response of mice as predicted by the top 9 models from the regression analysis.** The top 9 models together reflect $76.2\%$ of all selected models. Median reticulocyte response in the first phase (solid lines) and second (dashed lines) phase, as predicted from the 9 most-frequently selected models (individual panels). .The phase is defined with respect to the breakpoint (inset), such the first phase is the period before the breakpoint, the second phase the period after. Percentages in the titles of (A-P) reflect the frequency with which each model was selected. Line color indicates parasite nutrient (pABA) treatment.

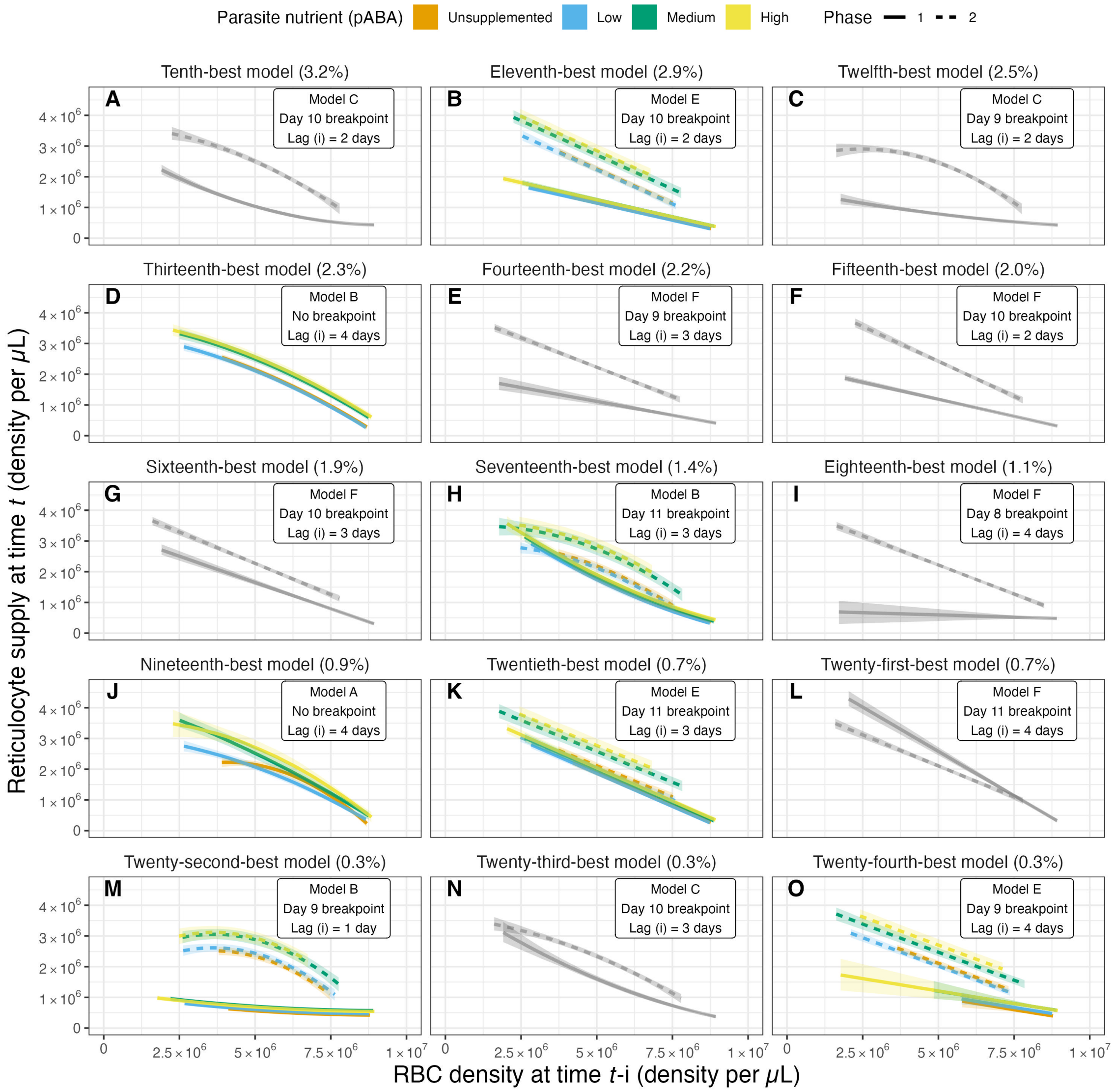


**Figure S6. The reticulocyte supply response of mice as predicted by the tenth-twenty-fourth most frequently selected models.** The top 24 models together reflect $98.9\%$ of all selected models. Median reticulocyte response in the first phase (solid lines) and second (dashed lines) phase, as predicted from each of the 9 most-frequently selected models (individual panels). The phase is defined with respect to the breakpoint (inset), such the first phase is the period before the breakpoint, the second phase the period after. Percentages in the titles of (A–P) reflect the frequency with which each model was selected indicate parasite nutrient (pABA) treatment; Panels with grey series display predictions from models that did not include pABA as an effect; the remainder of the panels display predictions from models that include pABA and the supply dynamics in each pABA treatment are displayed in different colors.

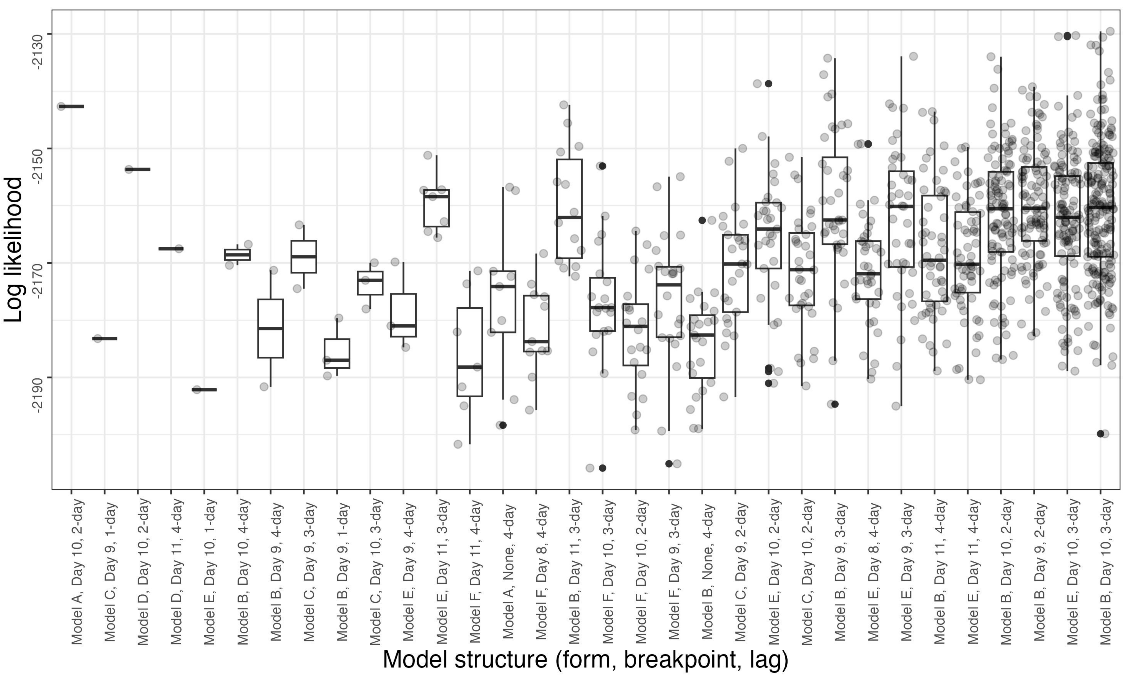


**Figure S7. Log likelihood distributions for all chosen models from the reticulocyte response regression analysis.** All models that were selected at least once in the reticulocyte response analysis are displayed. Models are arranged according to the frequency with which they were selected, i.e., with the model selected least often on the left and most often on the right.

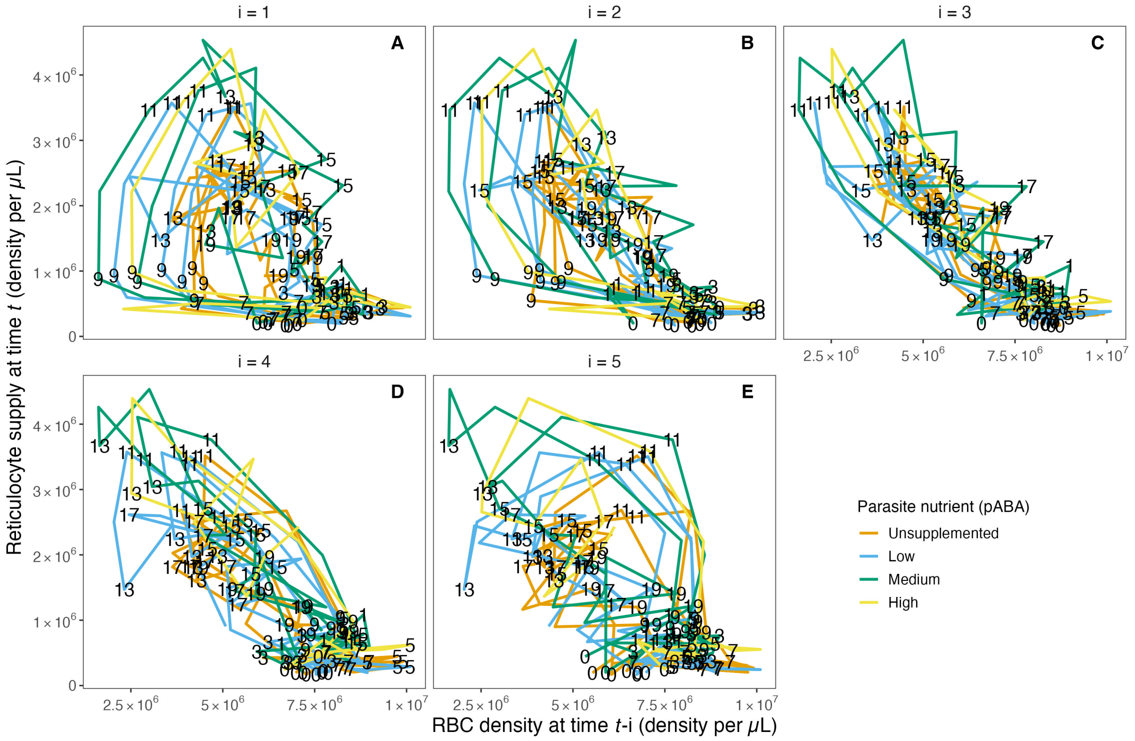


**Figure S8. Relationship between reticulocyte supply and lagged RBC (red blood cell) density in the raw data.** The relationship between raw RBC density on day $t-1$ (A), $t-2$ (B), $t-3$ (C), $t-4$ (D) and $t-5$ (E) and raw reticulocyte density on day $t$ for each of the four parasite nutrient (pABA) treatments. The paths are labelled with time $t$, i.e., a point on the path in (A) labelled 8 shows the reticulocyte supply on day 8, given the RBC density on day 7. A point on the path in (B) labelled 8 shows the reticulocyte supply on day 8 given the RBC density on day 6. Panels (C-E) work analogously but for RBC densities lagged by 3, 4 and 5 days.

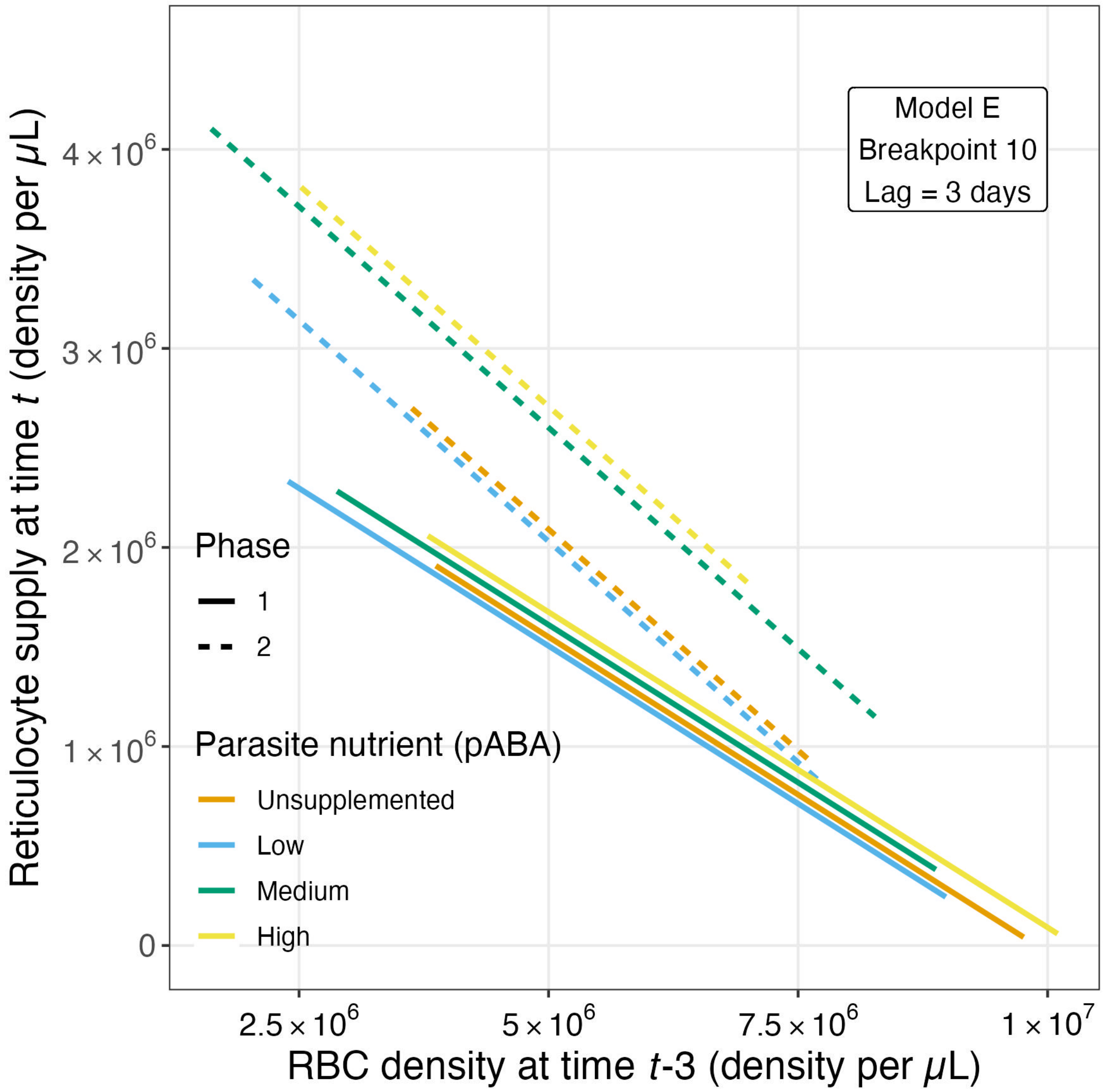


**Figure S9. The reticulocyte supply response as predicted from the best model yielded in an analysis of the raw data.** The top model was of model form Model E (Table 4, *Main text*) with a breakpoint at day 10 post-infection and specified a reticulocyte response lag of 3 days. Colors reflect parasite nutrient (pABA) treatment.

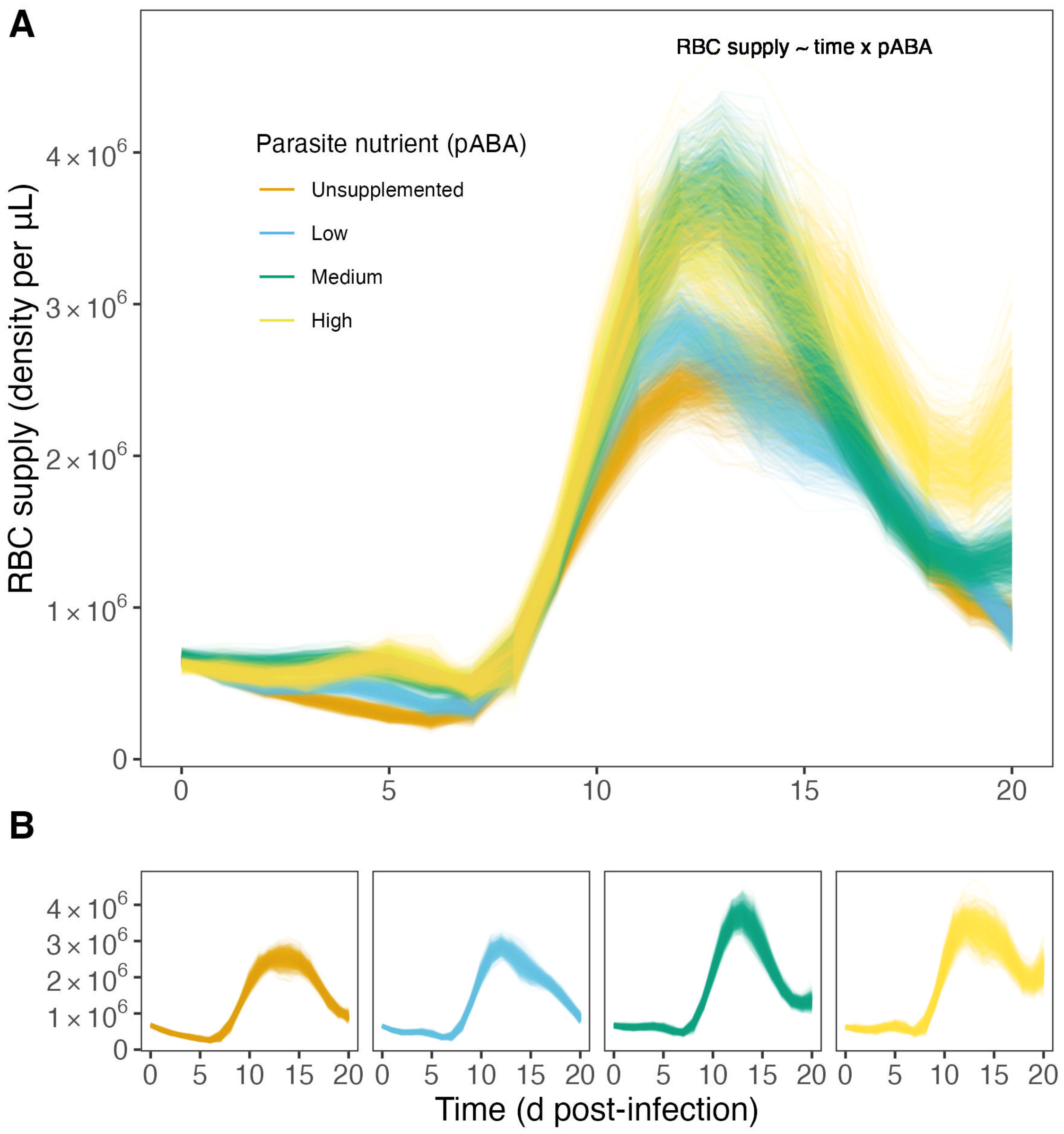


**Figure S10. The dynamics of reticulocyte supply during malaria infeciton, as predicted from the generalized additive model analysis.** (A) Predicted reticulocyte supply dynamics according to the top performing model (Model A, Table S1, *Supplementary Material*). Each line represents an iteration from the regression analysis. (B) As in A, but parsed by parasite nutrient treatment (i.e., pABA concentration) for ease of interpretation.
